# Microglia promote neurodegeneration and hyperkatifeia during withdrawal and prolonged abstinence from binge alcohol

**DOI:** 10.1101/2024.11.26.625461

**Authors:** Elizabeth McNair, Lamar Dawkins, Baylee Materia, Grace Ross, Alexandra Barnett, Puja Nakkala, Liya Qin, Jian Zou, Viktoriya Nikolova, Sheryl Moy, Leon G. Coleman

## Abstract

Proinflammatory microglial polarization, neuronal death, and hyperkatifeia/negative affect during withdrawal are key features of alcohol use disorder (AUD). However, the role microglia play in the development of AUD-related neuronal and behavioral pathology is unclear. Given the ability of microglia to regulate neuronal function, it was hypothesized that proinflammatory microglia promote neuronal death and hyperkatifeia during prolonged abstinence from binge alcohol. Proinflammatory signaling and affective state were assessed in mice either during acute withdrawal (24h) or abstinence (>4 weeks) to binge alcohol exposure. Ten days of binge alcohol increased proinflammatory gene signaling 24h after EtOH, which lasted weeks into withdrawal. Alcohol reduced brain-derived neurotrophic factor (BDNF) in hyperkatifeia-associated regions (i.e., the central amygdala and infralimbic cortex) during acute withdrawal and caused persistent microglial structural changes and loss of microglial BDNF in the BNST during abstinence. This was associated with increased anxiety-like behavior and hyperarousal, with persistent enhancement of conditioned fear memory during abstinence. Inhibition of proinflammatory microglia with Gi designer receptors exclusively activated by designer drugs (DREADDs) blocked neuronal death and prevented persistent proinflammatory gene induction and hyperkatifeia in female mice. Thus, this identifies a direct role for microglia in the development of AUD-related neuropathology and behavioral dysfunction, implicating microglia as cellular targets for the prevention of AUD phenotypes.

## Introduction

Nearly 95 million individuals suffer from alcohol use disorder (AUD) worldwide^1^. AUD is characterized by difficulty in ceasing alcohol use despite adverse effects on health and/or social life^2^. Current treatments for AUD include pharmacotherapies (naltrexone, acamprosate, and disulfiram), behavioral interventions, and support groups^3,4^. Despite these options, ∼60% of individuals with AUD relapse within the first year after treatment^5,6^. Various health and social factors promote AUD progression, maintenance and relapse, with persistent stress or negative affective states, also known as hyperkatifeia^7,8^, emerging as a significant contributor^9–14^. Alcohol (i.e., ethanol) misuse is also highly co-morbid with mood and affective disorders such as depression and post-traumatic stress disorder (PTSD)^15–17^. However, no current therapies for AUD target these persistent alcohol-induced hyperkatifeia states. Therefore, the high rates of AUD prevalence, difficulty to effectively treat AUD, and the association of AUD with affective dysfunction warrant a deeper understanding of alcohol-induced hyperkatifeia.

The natural history of AUD involves stressful life experiences followed by the initiation of binge drinking episodes^18–21^. These experiences interact with more proximal stressors to produce alcohol-seeking behavior^22^, with stress caused by repeated cycles of binge alcohol use and withdrawal synergizing with lifetime stressors to promote hyperkatifeia^23^. Therefore, treatment strategies that interrupt this cycle by diminishing alcohol-induced hyperkatifeia may be effective at reducing relapse or preventing AUD-related mood disorders. Several studies have focused on neuronal activity in key brain regions and circuits associated with alcohol-induced hyperkatifeia and anxiety-like behavior, such as the bed nucleus of the stria terminalis (BNST), central amygdala (CeA), and infralimbic cortex (IL). The BNST has been tied to aversive states and dysphoria during drug withdrawal^24–26^, as well as conditioned and unconditioned anxiety responses^27^. The CeA plays a role in the formation of anxiety-like behaviors with alcohol dependence, such as fear memory acquisition and extinction^28,29^, while the infralimbic cortex (IL) regulates stress-responses and subsequent alcohol-seeking behavior^30,31^.

Though much work has been done regarding the role of neurotransmission in these regions as it pertains to hyperkatifeia, relatively little is known regarding the role of glia and proinflammatory signaling in these regions. Proinflammatory cytokines are associated with stress and anxiety-like behavior^32,33^, while brain-derived neurotrophic factor (BDNF) signaling, an anti-inflammatory/trophic mediator, reduces alcohol-induced anxiety-like behavior in stress regions^34,35^. However, it is unclear is persistent proinflammatory signaling induced by alcohol promotes the development of long-lasting hyperkatifeia.

Proinflammatory signaling is a regulator of neurotransmission, particularly in disease states. Glia, such as microglia and astrocytes, impact neuronal firing, memory, and survival particularly when activated^36–38^. Prior reports indicate that binge ethanol induces proinflammatory polarization of microglia^39–41^. This involves the induction of innate immune pathways, such as nuclear transcriptional Nuclear factor kappa-light-chain-enhancer of activated B cells (NF-kB), with resulting expression of pro-inflammatory cytokines such as tumor necrosis factor alpha (TNFα), interleukin 1-beta (IL-1β), and type I interferons (IFNs)^42,43^. Correlations between proinflammatory gene expression and lifetime consumption of alcohol in postmortem human AUD brain suggest proinflammatory signaling increases with the number of exposures^44–47^. Proinflammatory stimuli, such as toll-like receptor agonists, promote alcohol drinking in rodents^48,49^, and microglial depletion prevents dependence-induced escalation in drinking^50,51^, suggesting a functional role for proinflammatory microglia in AUD. Further, neurodegeneration of the orbitofrontal cortex (OFC) is found in AUD. The OFC is essential for the extinction of well learned behaviors^52^, and OFC degeneration predicts persistent negative affect and relapse in AUD^53^. Microglia secrete soluble mediators that cause neuronal death in response to alcohol *in vitro*^45,47^. However, it is unknown if proinflammatory microglia promote neuronal death in the OFC *in vivo,* which could contribute to the inability to extinguish persistent hyperkatifeia in AUD.

In this study, alcohol-induced proinflammatory signaling, hyperkatifeia, and OFC neurodegeneration were assessed after two binge ethanol paradigms (4 or 10 consecutive daily binges). Proinflammatory vs trophic balance in stress-regulating regions involved in alcohol-induced hyperkatifeia were measured, in addition to anxiety-like behavior (thigmotaxis), hyperarousal (startle response), and extinction of conditioned fear memory (a PTSD-like phenotype and measure of behavioral flexibility^54,55^) during abstinence. To determine the role of microglia, microglia-specific Gi coupled designer receptors exclusively activated by designer drugs (DREADDs) were used that previously were reported to blunt microglial proinflammatory responses to alcohol^56–58^. This allowed for the specific assessment of the role of proinflammatory microglia in alcohol-induced OFC neurodegeneration and persistent hyperkatifeia.

## Materials and Methods

### Animal models

Adult 8-week-old C57BL/6J wild type (WT) mice were purchased from the Jackson Laboratory (Bar Harbor, Maine, Stock # 000664). Mice were given 1 week of habituation prior to alcohol treatment. For Gi DREADD inhibition, *CX3CR1.Cre^ERT2^.hM4di* mice were bred in-house from *CX3CR1.Cre^ERT2^* (JAX strain 020940, 129P2 background) and DIO.hm4di mice (JAX strain 026219, 129S background) obtained from Jackson Labs. Pups were weaned at 28 days old and housed with same-sex littermates. Protocols were in accordance with NIH regulations (Protocols 23-230.0 and 23-149.0) and approved by the University of North Carolina at Chapel Hill Institutional Animal Care and Use Committee (IACUC).

### Repeated binge ethanol treatment

Adult WT mice were given daily intragastric (i.g.) doses of isocaloric maltose dextran (9g/kg)^59^ or alcohol (i.e., ethanol, 5g/kg/d) for either 4 or 10 consecutive days to model human binge drinking. This dose was chosen as it produces blood alcohol concentrations over a 24-hour period equivalent to a binge-level dose in humans of 1.4g/kg (AUC: 230.7mM x hour, ∼5-7 standard drinks)^60^. No differences in weight were found between treatment groups (Figure 1B). Mice were anesthetized with isoflurane and sacrificed by transcardial perfusion with saline, either 24 hours (acute withdrawal) or 5 weeks into abstinence following behavioral testing. For acute withdrawal, two separate treatment cohorts were performed: Cohort 1 – Control N= 4 male, 5 female; EtOH N= 3 male, Cohort 2 – Control N=7 males, 6 females; EtOH N=8 males, 5 females.

**Figure 1:**
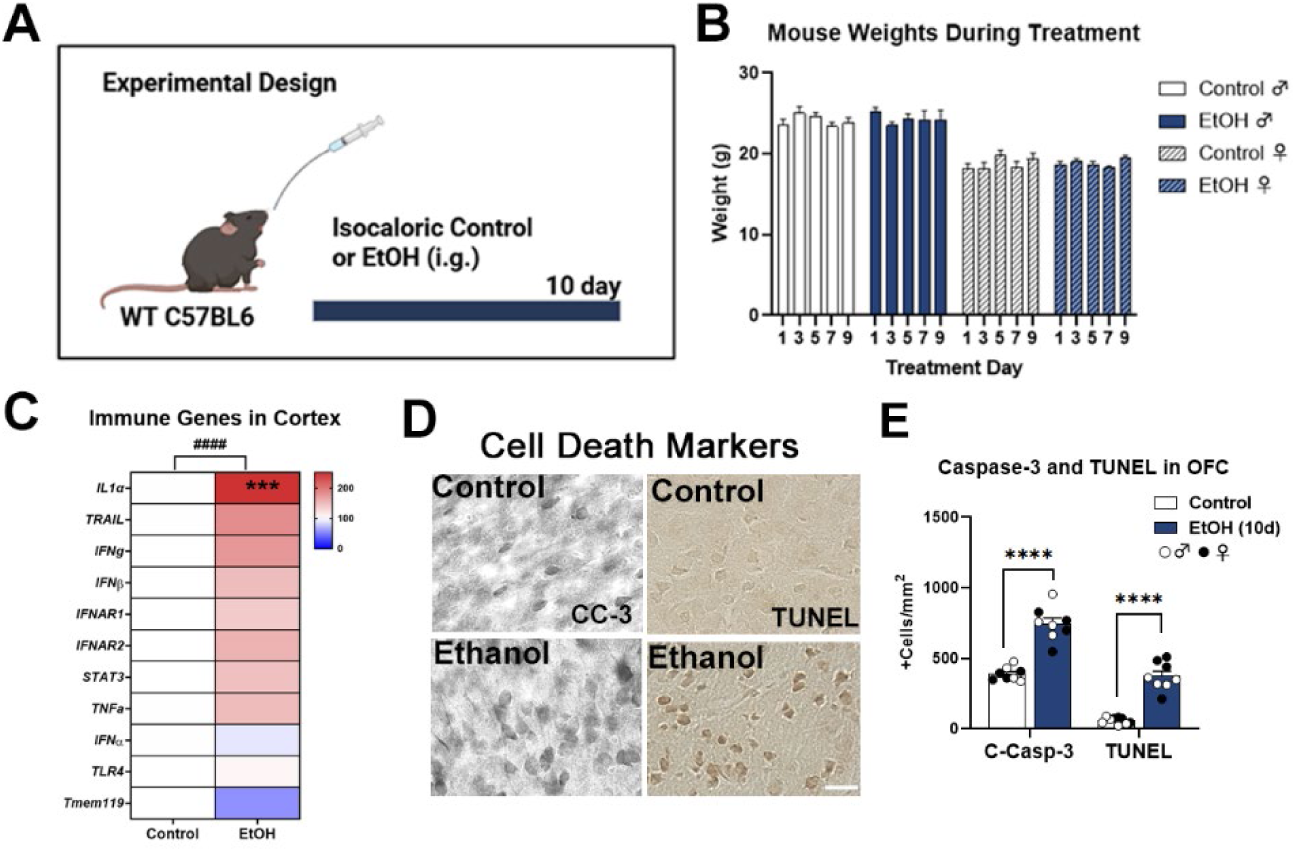
Cortical proinflammatory gene induction and neurodegeneration with binge alcohol. **(A)** Mice received isocaloric maltose dextran (9g/kg/d, i.g.) or binge ethanol treatment (EtOH, 5g/kg/d, i.g.) for 10 days with assessments 24 hours after the last treatment. Sexes were combined when no sex effects were observed. **(B)** No change in mouse weight was found across the treatment. **(C)** Ethanol induced proinflammatory gene expression in cortex of both males and females at 10 days of ethanol. 2-way ANOVA main effect of ethanol: F_1,150_=24.8, ####*p<*0.0001, \*\*\**p<*0.001 Sidak’s post-test. Control N=5 male, 5 female; EtOH N=4 male, 4 female. **(D)** Representative images of increased cleaved caspase-3 (CC-3, C-Casp-3) and TUNEL stain in the OFC. Scale bar: 40µm. **(E)** Quantification of CC-3 and TUNEL cell death stain in the OFC finding ∼2-fold increase in each cell death marker in both males and females. Males-open circles, females-black circles. Control N=5M, 5F; 10-day ethanol N=5M, 4F. \*\*\*\**p<*0.0001, *t-*tests.

### Organotypic brain slice culture

Organotypic brain slice cultures (OBSCs) were generated from *CX3CR1.Cre^ERT2^.hM4di* mice using our standard protocols^56,57,61^ with approval of the UNC Institutional Animal Care Use Committee (IACUC Protocol 24–096). OBSCs were prepared from male and female P7 mouse brain, as previously reported^56,57,62^. Pups were sacrificed by decapitation. The hippocampal-entorhinal complex was dissected while in Gey’s buffer (Sigma-Aldrich, St. Louis, MO). Slices were then prepared by transversely cutting the regions with McIlwain tissue chopper (375 μm thickness), and placed onto 30 mm diameter Millicell low height culture membrane inserts (Millipore, PICMORG50) , with 10–13 slices/ insert (i.e., culture well) as reported previously^40,42,45,47,63–68^. This requires ∼8-10 pups per preparation. Slices were maintained in MEM containing HEPES (25 mM) and Hank’s salts, with 25% horse serum (HS), 30mM glucose, and 2 mM L-glutamine for 7 days in vitro, followed by 4DIV in MEM + 12% HS, and then MEM + 6% HS. To induce Cre recombination, 4-hydroxy tamoxifen (4-OH-TAM, 10ng/mL) was added 24 hours prior to experimentation. For ethanol treatment, slices were treated for 96 hours followed by a 24-hour withdrawal period previously reported to result in alcohol withdrawal-induced cell death^45,47^. Cell death was measured with propidium iodide (PI) labeling, as done previously^45,47,57^. PI-labeled images were quantified using ImageJ (v 1.53c). The area of fluorescently labeled PI per total section area was measured. Images are presented as color inverted for ease of visualization. Each data point was an individually assessed brain slice.

### Microglial Gi DREADD inhibition *in vivo*

Heterozygous male and female *CX3CR1.Cre^ERT2^.hM4di* mice were given tamoxifen (75 mg/kg/d, i.p.) once daily for 5 days at 12 weeks of age to induce recombination. Mice were then given 4 weeks for repopulation of peripheral monocytes to minimize peripheral hM4di expression prior to treatment with water or ethanol (5g/kg/d, i.g. for 10 days) +/-clozapine N-oxide (CNO,3mg/kg, 10 hours after ethanol). Mice were sacrificed by transcardial perfusion 24 hours after the final dose or after 5 weeks of behavioral testing.

### Perfusion and tissue collection

Mice were sacrificed by transcardial perfusion with cold 0.1 M phosphate-buffered saline (PBS). Brains were removed from the skull and hemisected or taken whole. If hemisected, one hemisphere was dissected for cortex and hippocampus, then snap-frozen in liquid nitrogen for RNA analysis. The contralateral hemisphere was drop-fixed in paraformaldehyde (PFA, 4%) for immunohistochemistry (IHC). Brains were dehydrated in 30% sucrose and then coronal sections were made using a sliding microtome (40 μm, MICROM HM450; ThermoScientific, Austin, TX). Serial sections were placed in 12-well plates and stored in cryoprotectant (30% glycol/30% ethylene glycol in PBS) at −20°C.

### mRNA isolation and RT-PCR

mRNA was isolated from whole brain, frozen cortex, or hippocampal tissue consistent with previous reporting^69^. Samples were homogenized with Trizol (Invitrogen), and RNA was isolated by chloroform extraction. The nanodrop 2000^TM^ spectrophotometer was used to quantify RNA concentration and reverse transcription was completed with 10μL of diluted RNA and 40μL of reverse transcription mix. SYBR green PCR master mix (Applied Biosystems; 4368702) was used to complete RT-PCR. Primer sequences were found using the Harvard primer database^70^ or designed de-novo using the National Library of Medicine Primer-BLAST tool. Primer sequences are in Table 1. Housekeeping genes 18S and β-actin were used to normalize expression of genes of interest using the cycle threshold (Ct) values obtained for each product. The ΔΔCt method was used to compare relative differences between groups and the percent change relative to 18S was used in analysis as reported previously^45,63,65^.

### Immunohistochemistry (IHC), immunofluorescence (IF), and TUNEL staining

IHC and IF were performed using our published protocols^45,63,65,69^. Briefly, for IHC free-floating tissue sections (40 μm) were washed in 0.1M PBS, quenched in 0.6% H_2_O_2_ for 30 minutes to inhibit endogenous peroxidases, then incubated for 1 hour at 70°C in pH=6.0 1X Antigen Retrieval Citra solution (Fisher, Pittsburgh PA; NC9935936). For membrane permeability, sections were blocked at room temperature for 1 hour in 4% normal serum with 0.1% Triton X-100 (Millipore Sigma, Burlington Massachusetts; T8787). Sections were incubated in primary antibody in blocking solution overnight at 4°C. Primary antibodies are in Table 2. Sections were washed in PBS and incubated in the corresponding biotinylated secondary antibody (1:200, Vector Laboratories, Newark CA) for 1 hour at room temperature. After additional washes, sections were incubated in avidin-biotin complex (Vector Vectastain ABC Kit; PK-6100) for 1 hour. 3,3’-Diaminobenzidine tetrahydrochloride hydrate (Sigma-Aldrich; D4293-50SET) was used for visualization. Sections were mounted on glass slides, air dried, dehydrated in a series of ethanol concentrations, and covered with Cytoseal (Fisher; 22-050-262). For IF, H_2_O_2_ incubation was omitted, and after primary antibody incubation sections were washed and incubated at room temperature in the appropriate Alexa Fluor conjugated secondary antibody (1:1000; Invitrogen). Sections were washed and mounted, allowed to dry, and cover slipped with Prolong Gold Anti-Fade mounting medium with DAPI (ThermoFisher; P36971). Terminal deoxynucleotidyl transferase dUTP nick end labeling (TUNEL) staining was performed using the Click-iT^TM^ TUNEL Colorimetric IHC Detection kit (ThermoFisher) according to the manufacturer’s instructions.

Representative images of the tissue were taken on the Keyence BZ-X800 Microscope and its accompanying BZ-X00 Analyzer software (v. 1.1.1.8) was used for microscopic analysis. For each region of interest, images were taken of ∼3 identical sections per subject, selected by desired bregma according to the mouse brain atlas (Paxinos and Watson). By regions, bregma were: infralimbic cortex (IL, 1.98mm), anterior cingulate and prelimbic cortex (2.10 mm), orbital frontal cortex (OFC, 2.10mm), dorsal and ventral bed nucleus of the stria terminalis (BNST, 0.14mm), central amygdala (CeA, −1.82mm), and basal lateral amygdala (BLA, −1.70mm). Positive staining was reported in immunoreactive (+IR) pixels/mm^2^ area of the region assessed. For colocalization, the number of target pixels within the larger cell area (i.e., BDNF puncta within Iba-1+ microglia was measured as reported^71^) using the BZ-X00 Analyzer software. Confocal z-stacks were taken with a 1µm interval step size on an ECHO spinning disk confocal orthogonal images made using ImageJ^TM^ to confirm co-localization (Supplemental Figure 1).

### Behavioral assessments

Behavioral testing was performed from days 3-31 during ethanol withdrawal and forced abstinence. Mice underwent repeated testing that was ordered from least stressful to most stressful to limit carry-over effects and were given extended recovery time between each behavioral assay as reported previously^72–74^.

#### 2.8.1 Open field

Exploratory activity in a novel environment was assessed by a one-hour trial in an open field chamber (41 cm x 41 cm x 30 cm) crossed by a grid of photobeams (VersaMax system, AccuScan Instruments) as reported^75,76^. The number of photobeams broken during the trial was counted, in addition to measurements for locomotor activity (total distance traveled) and vertical rearing movements. Anxiety-like behavior was indexed by time spent in the center region.

#### 2.8.2 Acoustic startle test

This procedure was used to assess auditory function, reactivity to environmental stimuli, and sensorimotor gating. The test was based on the reflexive whole-body flinch, or startle response, following exposure to a sudden noise. Measures were taken of startle magnitude and prepulse inhibition, which occurs when a weak pre-stimulus leads to a reduced startle in response to a subsequent louder noise (San Diego Instruments SR-Lab system). Mice were placed into individual small Plexiglas cylinders within larger, sound-attenuating chambers. Piezoelectric transducers underneath each cylinder recorded the force of startle responses. The chambers included a ceiling light, fan, and a loudspeaker for the acoustic stimuli. Each session began with a five-min habituation period, followed by 42 trials (7 of each type): no-stimulus trials, trials with the acoustic startle (40 msec; 120 dB) alone, and trials with a prepulse stimulus (20 msec; either 74, 78, 82, 86, or 90 dB) occurred 100 msec before the onset of the startle stimulus. Measures were taken of the startle amplitude for each trial across a 65-msec sampling window, across levels of prepulse.

#### 2.8.3 Morris water maze

The water maze was used to assess spatial and reversal learning, swimming ability, and vision as reported^75^. The water maze consisted of a large circular pool (diameter = 122 cm) partially filled with water (45 cm deep, 24-26° C), located in a room with numerous visual cues. The procedure involved two 5-day phases: acquisition in the hidden platform task, and a test for reversal learning (an index of cognitive flexibility). In both phases, mice were tested for their ability to find a submerged, hidden escape platform (diameter = 12 cm). For each trial, the mouse was placed in the pool at 1 of 4 possible locations (randomly ordered) and then given 60 sec to find the platform. If the mouse found the platform, the trial ended, and the animal was allowed to remain 10 sec on the platform before the next trial began. If the platform was not found, the mouse was placed on the platform for 10 sec and then given the next trial. Measures were taken of latency to find the platform and swimming speed via an automated tracking system (Noldus Ethovision). 2-3 hours after the final trial on day 5, mice were given a 1-min probe trial in the pool with the platform removed. Selective quadrant search was evaluated by measuring percent time in the quadrant where the platform (the target) had been located during training and number of crosses over the platform area, versus the corresponding areas in the other three quadrants. Following the acquisition phase, mice were tested for reversal learning, using the same procedure as described above. In this phase, the hidden platform was re-located to the opposite quadrant in the pool. As before, measures were taken of latency to find the platform. On day 5 of testing, the platform was removed from the pool, and the group was given a probe trial to evaluate reversal learning.

#### 2.8.4 Conditioned fear with extinction

This standard test for learning, memory, and extinction was conducted across 3 days, using a Near-Infrared image tracking system (MED Associates, Burlington, VT). On the first day, mice were given a 7-min training session. Mice were placed in the test chamber, contained in a sound-attenuating box, and allowed to explore for 2 min. The mice were then exposed to a 30-sec tone (80 dB), which co-terminated with a 2-sec scrambled foot shock (0.4 mA). Mice received 2 additional shock-tone pairings during the training session. Context-dependent learning was evaluated on the second day of testing. Mice were placed back into the original test chamber, and levels of freezing (immobility) were determined across a 5-min session. On the third day of testing, mice were evaluated for associative learning to the auditory cue and extinction of freezing responses. To conceal contextual cues, the conditioning chambers were modified using a Plexiglas insert to change the wall and floor surface, and a mild novel odor (70% ethanol) was added to the sound-attenuating box. Mice were placed in the modified chamber and allowed to explore. After 2 min, mice were given 50 presentations of the 30-sec tone, with 5 sec between each presentation. Levels of freezing before and during the stimuli were obtained by the image tracking system.

### Statistical analyses

Statistical tests were performed in GraphPad PrismTM version 10.4. t-tests were used for preplanned orthogonal contrasts between two groups. For assessments comparing multiple groups with one categorical independent variable, 1-way ANOVAs were used. For comparisons with two factors 2-way ANOVAs were performed with or without repeated measures as appropriate. Sidak’s or Dunnett’s multiple comparisons tests were used as appropriate. Outliers were removed according to Grubb’s test.

## Results

### Binge ethanol acutely induces proinflammatory signaling and cortical neuronal death

Studies in postmortem human brain suggest the extent of proinflammatory induction is related to the lifetime duration of alcohol use^43,77^. To test this directly, proinflammatory signaling and neuronal death were assessed in mice after 10 daily consecutive alcohol binges. Mice received 5g/kg/d by gavage to model human binge drinking with sacrifice 24 hours after the last dose of alcohol as reported previously^47^ (Figure 1A). This model reproduces many of the neuroinflammatory findings seen in human AUD^47,78,79^, with modest proinflammatory induction in the liver (TNFα with no changes in IL-1β) and no increase in serum TNFα or IL-1β^80^. Mice do not show signs of severe withdrawal such as seizures and no changes in weight were found across the treatment (Figure 1B) and no signs of dehydration were observed by DLAM certified veterinary staff. Mice metabolize alcohol (i.e., ethanol) up to 10X faster than humans^60^. Thus, in mice, this dose results in a blood ethanol concentration AUC x time of 230.7mM x hour, which is seen in humans with ∼1.4g/kg or ∼7 standard drinks (14g standard drink, 70kg body weight)^60^. Twenty-four hours after ten alcohol binges, changes in proinflammatory gene expression were seen in the cortex with similar changes in males and females. A significant main effect of alcohol treatment was found (Figure 1C, F_1,150_=24.8, *####p*<0.0001) with a statistically significant increase in interleukin 1-alpha (*IL-1α*, \*\*\**p<*0.0001, Sidak’s post-test). Accompanying changes in proinflammatory gene expression in cortex were increased measures of neuronal death. In the OFC robust increases in the executioner caspase cleaved capsase-3 (CC-3) and TUNEL were seen after 10 ethanol binges (Figure 1D-E, \*\*\*\**p<*0.0001, t-tests). CC-3 was also increased in the PrL and IL, though TUNEL+ staining was not observed (data not shown). Caspase-3 cleavage does not always result in cell death and may suggest heightened vulnerability of the OFC consistent with a prior report^81^. Next, microglial structure and trophic support was assessed in cortical and subcortical regions after binge ethanol.

Microglial number as well as cell body and process area were assessed during early withdrawal in two treatment cohorts (24h after the last dose), as these measures have been used previously by many studies as surrogates for alterations in microglial phenotype that are complementary to gene expression assessments^47,82–84^. Microglial trophic support was assessed by immunofluorescence (IF) for Iba-1 and BDNF in the second cohort of mice (due to limited sections remaining in cohort 1) in regions associated with alcohol-induced stress and anxiety^85^ including the dorsal bed nucleus of the stria terminalis (dBNST), vBNST, central amygdala (CeA), and infralimbic cortex (IL). In the dBNST alcohol had no effect on microglial number; however, a main effect of treatment was found on microglial cell body and process area that reached statistical significance in males (Figure 2A-C, F_1,38_=9.2, *##p=*0.0043, *\*p<*0.05, Sidak’s post-test) along with a significant effect of sex on microglial area (F_1,38_=22.17, *p<*0.0001). This increase in microglial cell body and process area is consistent with a hyper-ramified state observed previously with alcohol^82,86^. No differences were found in total or microglial BDNF in the dBNST (Figure 2D-G). However, there was a notable sex difference in total (F_1,23_=19.36, *p=*0.0002) and microglial BDNF (F_1,23_=27.4, *p<*0.0001) found similar to other brain regions. The vBNST showed a similar pattern, with no change in microglial number. A main effect of alcohol was found on microglial cell body and process area that was significantly different in males (Figure 2H-J, F_1,39_=7.9, *##p=*0.0077, *\*p<*0.05, Sidak’s post-test). No changes in total BDNF were found in the vBNST, though a significant sex difference was found (Figure 2K-L, F_1,23_=9.7, *p=*0.005). A main effect of sex on microglial BDNF in the vBNST was also observed (F_1,23_=41.8, *p<*0.0001), with ethanol causing a trend toward a reduction that was driven by the male subjects (Figure 2M-N, F_1,23_=3.8, *p=*0.06).

**Figure 2:**
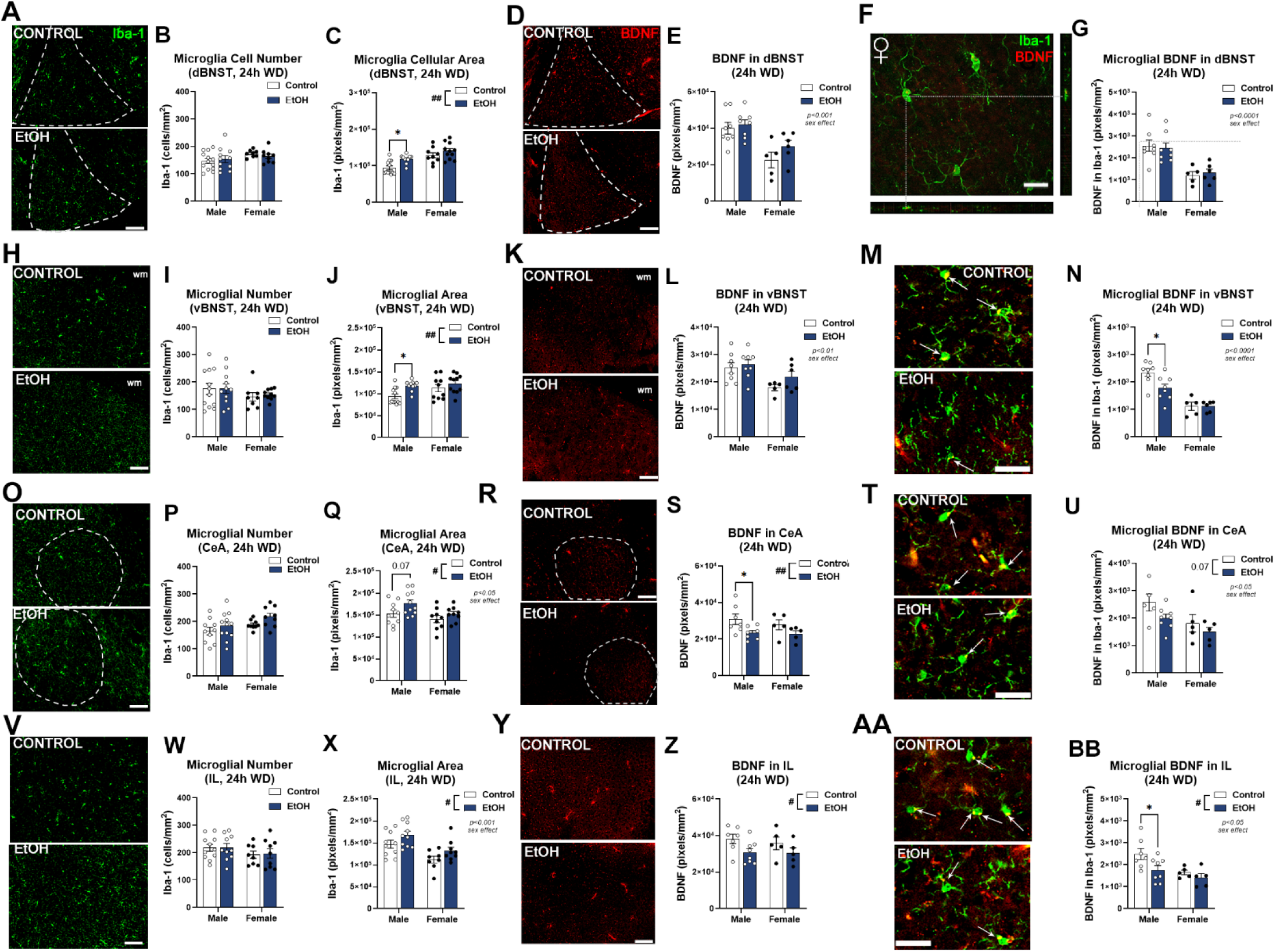
Microglial structure and BDNF in alcohol stress-related brain regions during early withdrawal from alcohol. Mice received isocaloric maltose dextran (9g/kg/d, i.g.) or binge ethanol treatment (EtOH, 5g/kg/d, i.g.) for 10 days with assessments 24 hours after the last treatment. Two treatment cohorts were performed with microglia structure measured on both cohorts and microglial BDNF measured on cohort 2. **(A)** Representative images of Iba-1 in the dBNST. **(B)** Ethanol had no impact on microglia cell number in the dBNST. Scale bar: 100µm **(C)** Ethanol altered dBNST microglia cell body and process area with a main effect of treatment and a significant reduction in males. 2-way ANOVA F_1,38_=9.2, *##p=*0.0043, *\*p<*0.05, Sidak’s post-test. A significant sex effect was also found F_1,38_=22.17, *p<*0.0001. **(D)** Representative images of total BDNF in the dBNST. Scale bar: 100µm **(E)** Ethanol had no effect on total BDNF in the dBNST, though a main effect of sex was found (F_1,23_=19.36, *p=*0.0002). **(F)** Orthogonal image of microglial BDNF in the dBNST confirming co-localization. Scale bar: 25µm. **(G)** Ethanol had no effect on microglial BDNF in dBNST, though a main effect of sex was found (F_1,23_=27.4, *p<*0.0001). **(H)** Representative images of Iba-1 in the vBNST. Scale bar: 100µm **(I)** Ethanol had no impact on microglia cell number in the vBNST **(J)** A main effect of ethanol on microglial cell body and process area was found in the vBNST that reached statistical significance in males. F_1,39_=7.9, *##p=*0.0077, *\*p<*0.05, Sidak’s post-test. **(K)** Representative images of total BDNF in the vBNST. Scale bar: 100µm. wm: white matter **(L)** Ethanol had no effect on total BDNF in the dBNST. A main effect of sex was observed. F_1,23_=9.7, *p=*0.005. **(M)** Representative image of microglial BDNF in the vBNST. Scale bar: 25um **(N)** A trend toward a main effect of ethanol on microglial BDNF was found in the vBNST driven primarily by the male subjects. F_1,23_=3.8, *p=*0.06. A main effect of sex was found F_1,23_=41.8, *p<*0.0001. *\*p<*0.05, Sidak’s post-test. **(O)** Representative images of Iba-1 in CeA. Scale bar: 100µm **(P)** Ethanol had no impact on microglia cell number in the CeA **(Q)** A main effect of ethanol was found on microglial cell body and process area (F_1,35_=4.9, *#p=*0.033) with a significant sex effect (F_1,35_=5.6, *#p=*0.02) **(R)** Representative images of total BDNF in the CeA. Scale bar: 100µm **(S)** A main of ethanol was found on BDNF in the CeA that reached a statistically significant reduction in males. F_1,20_=7.2, #*p=*0.01. *\*p<*0.05, Sidak’s post-test. **(T)** Representative image of microglial BDNF in the CeA. Scale bar: 25um **(U)** A trend toward a main of ethanol was found on microglial BDNF in the CeA (F_1,19_=3.6, *p=*0.07) with a main effect of sex (F_1,19_=7.3, *p*= 0.01). **(V)** Representative images of Iba-1 in the infralimbic cortex (IL). Scale bar: 100µm **(W)** Ethanol had no effect on microglial number in the IL. **(X)** Ethanol increased microglial cell body and process area. F_1,33_=5.8, *#p=*0.02. A significant sex effect was also found F_1,33_=17.2, *p=*0.0002. **(Y)** Representative images of BDNF in the IL. Scale bar: 100µm **(Z)** Ethanol caused a significant reduction in BDNF in the IL. F_1,21_=5.3, #*p=*0.03**. (AA)** Representative images of microglial BDNF in the IL. Scale bar: 25um **(BB)** Ethanol caused a significant reduction in microglial BDNF in the IL that reached statistical significance in male subjects. F_1,21_=7.0, #*p=*0.015, \**p<*0.05, Sidak’s post-test. Dashed white lines denote region of interest.

In the CeA, microglial number was unchanged (Figure 2O-P), while a main effect of ethanol on microglial cell body and process area was found (Figure 2Q, F_1,35_=4.9, *#p=*0.033). Ethanol caused a significant effect on total BDNF in the CeA, with a statistically significant reduction in males (Figure 2R-S, F_1,20_=7.2, #*p=*0.01. *\*p<*0.05, Sidak’s post-test). A main effect of sex was found on microglial BDNF in the CeA (F_1,19_=7.3, *p*= 0.01), with ethanol causing a trend toward a reduction in microglial BDNF in males (Figure 2T-U, F_1,19_=3.6, *p=*0.07). In the IL microglial numbers were unchanged, and a main effect of ethanol to increase microglial area was seen across both sexes (Figure 2V-X, F_1,33_=5.8, *#p=*0.02). A main effect for a reduction in total BDNF was also found (Figure 2Y-Z, F_1,21_=5.3, #*p=*0.03) as well as a reduction in microglial BDNF that was driven mainly by the male subjects (Figure 2AA-BB, F_1,21_=7.0, #*p=*0.015, \**p<*0.05, Sidak’s post-test). Together, this identifies microglial structural changes in alcohol-related stress regions early during withdrawal from repeated binge ethanol that are more pronounced in males. It also finds a reduction in trophic support as measured by BDNF (CeA and IL) and microglial BDNF (vBNST, CeA, and IL) that is also more pronounced in males. Given these neuroimmune changes in alcohol stress-related brain regions, we next assessed behavioral consequences of repeated binge ethanol.

### Binge alcohol enhances anxiety-like behavior with persistent fear memory deficits during abstinence

Due to the proinflammatory phenotype changes in stress-related regions after 10 days of binge ethanol, affective behavioral analyses were assessed next. Since negative affect in AUD during withdrawal and abstinence is thought to promote recurrent use in AUD^8^, behaviors were assessed 3-31 days after the last ethanol dose in male mice (Figure 3A). Open field and acoustic startle were performed to assess anxiety-like behavior and hyperarousal, which can promote drinking^7^. Mice traveled significantly less distance in the center of the open field with 10 but not 4 ethanol binges suggesting increased anxiety-like behavior (Figure 3B, F_2,39_=2.9, *p=*0.066, \**p<*0.05, Sidak’s post-test), though total center time was unchanged (Figure 3C). Ten ethanol binges reduced fine motor movement, with a trend toward a reduction after 4 binges (Figure 3D, F_2,31_=6.8, *p=*0.037, \*\**p<*0.01, Sidak’s post-test). With ten ethanol exposures, mice had increased startle magnitudes across different pre-pulse decibels consistent with hyperarousal (Figure 3E, F_2,287_=43.112, *p<*0.0001, *\*\*\*\*p*<0.0001, Dunnett’s post-test). Since negative affect and cognitive dysfunction in AUD persist in humans, the Morris Water Maze with reversal and conditioned fear memory acquisition with extinction were assessed as measures of behavioral inflexibility^54^. In the Morris Water Maze, mice exhibited no differences in learning (Figure 3F). However, during reversal learning trials, which require proper OFC function, ethanol blunted performance in a dose-dependent manner, with a significant delay with 10 ethanol binges (Figure 3G, F_1,63_=10.44 ##*p*=0.002, \**p<*0.05 Sidak’s post-test) and reduced time spent in the reversal learning quadrant during the probe trial (Figure 3H-I, F_6,164_=2.512, *p=*0.02, \*\**p*<0.01). In Pavlovian fear conditioning, ethanol had no impact on the acquisition of conditioned fear cue association learning (Figure 3J). However, mice undergoing 10 ethanol binges failed to extinguish the conditioned fear memory cue association, while their control and 4 ethanol binge counterparts showed clear memory extinction (Figure 3K) Thus, binge ethanol promotes an anxiety-like state with hyper-arousal, and a loss of cognitive and affective behavioral flexibility. Given these behavioral phenotypes and persistent proinflammatory/trophic imbalance with prolonged abstinence from binge ethanol, the specific role microglia play in the development of this persistent proinflammatory and negative-affective state was assessed next.

**Figure 3.**
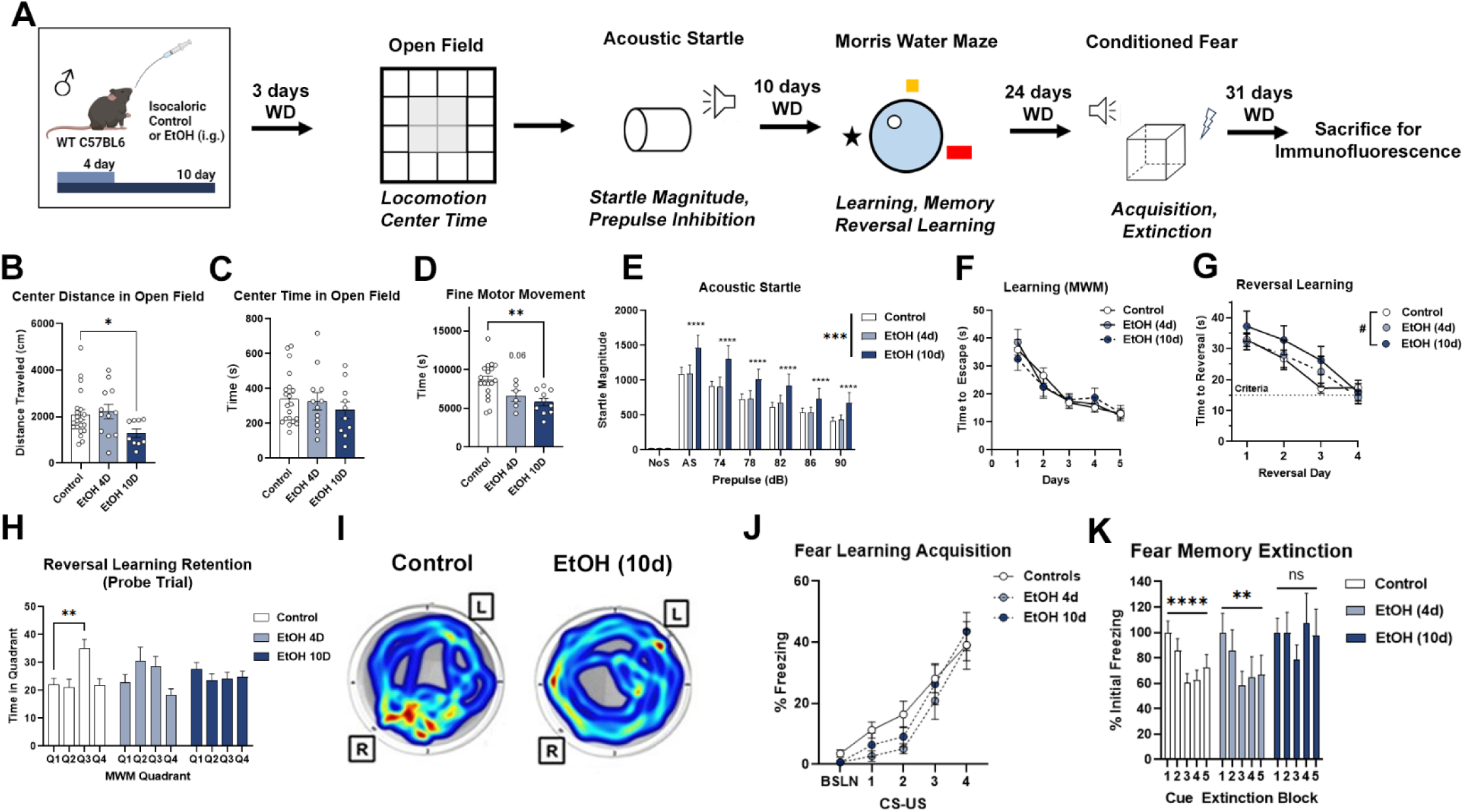
Negative affect and cognitive dysfunction with binge alcohol. **(A)** Mice underwent behavioral testing after treatment with isocaloric control (9g/kg/d, i.g.), no gavage control, or binge ethanol (EtOH, 5g/kg/d, i.g. for 4 or 10 days). At 3 days post-ethanol treatment, mice underwent open field testing for anxiety-like behavior, followed by acoustic startle for arousal. Morris water maze (MWM) testing began 10 days into abstinence for learning and reversal learning. Twenty-four days into withdrawal, the mice underwent conditioned fear testing for acquisition and extinction. Mice were sacrificed one week later for immunofluorescence. Isocaloric control and no gavage controls were combined due to lack of differences. **(B)** Mice receiving 10 days of binge ethanol traveled significantly less distance in the center of the open field (cm). 1-way ANOVA F_2,39_=2.9, *p=*0.066, *\*p<*0.05, Sidak’s multiple comparisons test. **(C)** No difference in time spent in the center across treatments. **(D)** 10 days of EtOH significantly reduced fine motor movement. 1-way ANOVA F_2,31_=6.8, *p=*0.037, \**\*p<*0.01, Sidak’s multiple comparisons test.. **(E)** Binge ethanol increased the magnitude of acoustic startle across pre-pulse intensities (dB) following 10 days of treatment. 2-way ANOVA_treatment_, F_2,287_=43.112, *p<*0.0001, *\*\*\*\*p*<0.0001, Dunnett’s post-test. **(F)** No change in learning with the Morris Water Maze was observed. Ten days of binge ethanol **(G)** increased latency during reversal days versus controls. 2-way ANOVA_treatment_ F_1,97_=4.08 #*p*<0.05 and **(H)** reduced time spent in the reversal quadrant (Q3) in EtOH treatment groups during probe trial after reversal learning 2-way ANOVA_tx x quadrant_ F_6,164_=2.512, *p=*0.02, \*\**p*<0.01, Dunnett’s post-test. **(I)** Representative heat map tracing during the reversal learning probe trial showing control mice spending a greater amount of time in the correct reversal quadrant than the ethanol-treated mice. R: correct reversal quadrant, L: initial learning quadrant **(J)** Ethanol had no effect on freezing acquisition in conditioned fear learning trials. **(K)** Control and 4-day binge ethanol groups showed extinction of conditioned fear memory, while mice receiving 10 ethanol binges showed no extinction. Control N=22, 4-day ethanol N=12, 10-day ethanol N=10. Male mice. \**p*<0.05, ns-no significant change, 1-way repeated measures ANOVA: Control F_2.76,58_=12.3, \*\*\*\**p*<0.0001; EtOH 4D F_2.3,25.4_=8.3, \*\**p=*0.0011; EtOH 10D F_1.8,16.0_=1.2, *p=*0.31.

### Lasting proinflammatory activation and loss of microglial BDNF with prolonged abstinence

Given the persistent negative affect after repeated binge ethanol and heightened proinflammatory response seen during acute withdrawal, microglial structure was assessed during prolonged abstinence following behavioral testing. Since ten ethanol binges caused lasting affective disruption, this rather that four ethanol binges was assessed. Male WT mice were assessed. After 31 days of abstinence from 10 days of binge ethanol, no changes were found in microglial number in the dBNST (Figure 4A-B). However, microglia showed persistent structural changes, evidenced by decreased microglia cell body and process area (Figure 4C, *\*p*<0.05) consistent with a dystrophic phenotype^87^. Though there was no change in total BDNF (Figure 4D-E), a significant reduction in microglial BDNF was found in the dBNST (Figure 4F-G, \**p<*0.05). In the vBNST, no change in microglial number was observed (Figure 4H-I), with a reduction in microglial cell body and process area found similar to the dBNST (Figure 4J, \**p<*0.05). Total BDNF was unchanged in the vBNST (Figure 4K-L), though a significant reduction in microglial BDNF was found (Figure 4M-N, \**p<*0.05). In the CeA, a slight increase in microglial number was seen with ethanol, though no robust changes in microglial area or microglial BDNF were found (Figure 4O-U). The IL likewise showed a slight increase in microglial number, with no persistent changes in neither microglial cell body and process area nor BDNF (Figure 4V-BB). Thus, persistent microglial structural changes and decreased microglial BDNF were found in the dBNST and vBNST. Given the role of BNST in stress phenotypes, these changes raised the question as to whether proinflammatory microglia promote persistent hyperkatifeia during abstinence.

**Figure 4.**
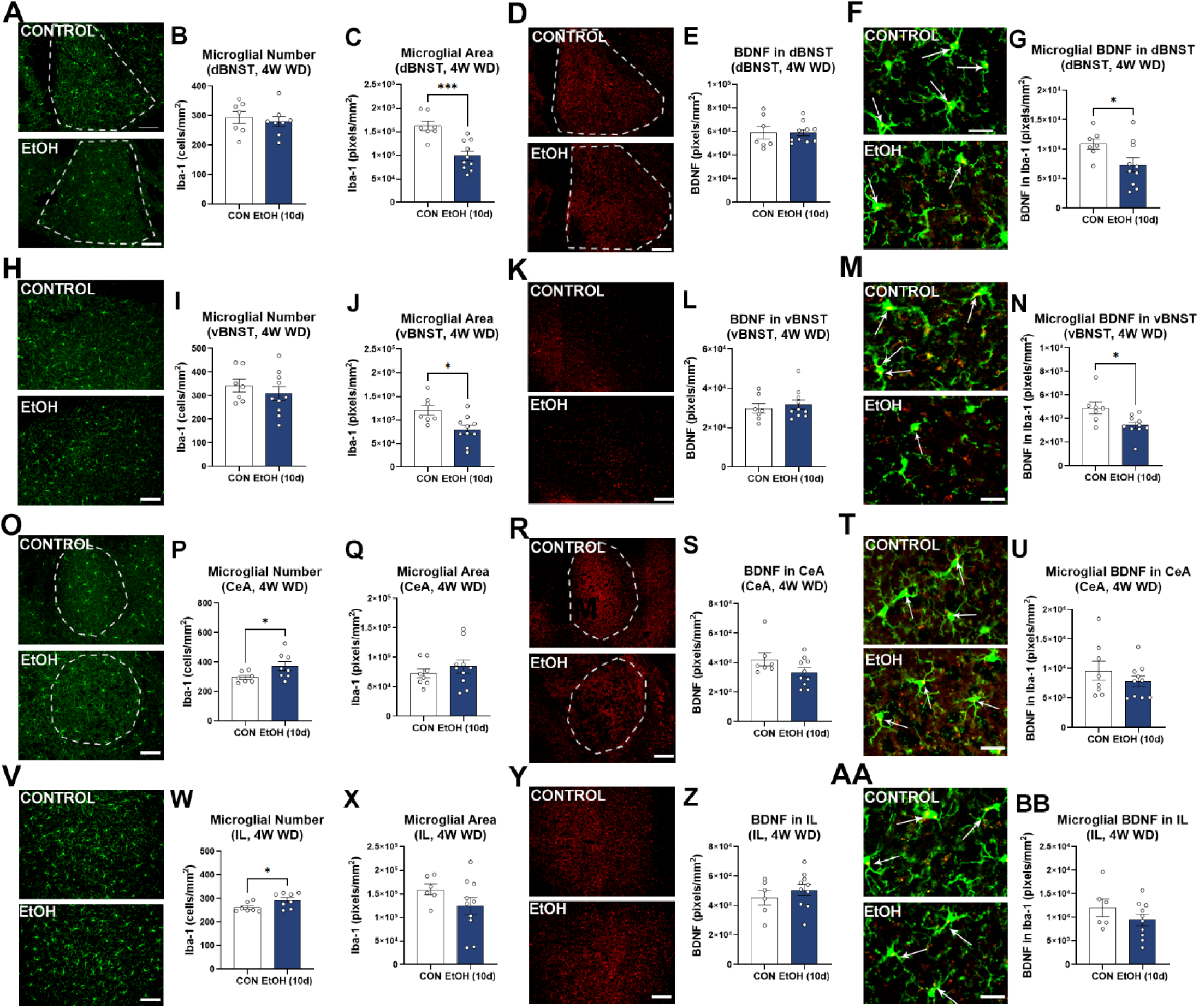
Persistent changes in microglia and BDNF with binge alcohol. IF assessments were performed four weeks into withdrawal after 10-day treatment with isocaloric control (9g/kg/d, i.g.) or binge alcohol (5g/kg/d, i.g.) and behavioral testing. N= Control=10 males, 10-day ethanol=10 males. **(A)** Representative images of Iba-1 in the dBNST. Scale bar 100µm. **(B)** No effect of ethanol on microglial number in dBNST. **(C)** Ethanol caused persistent reduction in the total area of microglial cell bodies and processes in the dBNST. \*\*\**p*<0.001, *t-*test. **(D-E)** Representative images and quantification for BDNF in the dBNST show no change 4 weeks into withdrawal. Dashed white lines denote region of interest. Scale bar 100µm **(F-G)** Representative image and quantification finding ethanol caused a persistent decrease in microglial BDNF in the dBNST four weeks into withdrawal. \**p*<0.05, *t-*test. Scale bar 25µm. Arrows denote colocalization of BDNF within microglia. **(H)** Representative images for Iba-1 in vBNST. Scale bar 100µm**. (I)** Ethanol had no effect on microglial number in the vBNST. **(J)** Ethanol caused a persistent decrease in microglial cell body and process area in the vBNST. *\*p*<0.05, *t-*test. **(K-L)** Representative image and quantification of BDNF in the vBNST finding no change. Scale bar 100µm**. (M-N)** Representative image and quantification finding a significant decrease in microglial BDNF in the vBNST four weeks into withdrawal. *\*p*<0.05, *t-*test. Scale bar 25µm. **(O)** Representative images for Iba-1 in the CeA. Scale bar 100µm**. (P)** Ethanol caused a slight increase in the number of microglia in the CeA. \**p<*0.05, *t-*test. **(Q)** Ethanol caused a trend toward an increase in total microglial area in the CeA. *p=*0.115, *t-*test. **(R-S)** Representative image and quantification of BDNF in the CeA finding a trend toward a reduction with ethanol. *p*=0.11, *t-*test. Scale bar 100µm**. (T-U)** Representative images and quantification finding no change in microglial BDNF in the CeA. **(V)** Representative image of microglia in the IL 4 weeks into withdrawal. **(W)** Ethanol caused a slight increase in the number of microglia in the IL. **(X)** Ethanol had no effect on total microglial area in the IL. *p=*0.19, *t-*test. Scale bar 25µm. **(Y-Z)** Representative images and quantification finding no change in BDNF in the IL. **(AA-BB)** Representative images and quantification finding no change in microglial BDNF in the IL. Dashed white lines denote region of interest

### Gi-induction in microglia prevents ethanol-induced neurodegeneration

Considering the co-occurring microglial proinflammatory polarization and neuronal death during acute withdrawal^47^, the ability of microglia to directly promote ethanol-induced neurodegeneration was assessed next. Microglia mobilize intracellular calcium stores to change their effector state^88^. Therefore, chemogenetic Gi DREADD (hM4di) activation was used to blunt microglial proinflammatory polarization as reported previously^58,63,65,89^. *CX3CR1.Cre^ERT2^* mice were bred with DIO.hM4di (DREADD) mice. Heterozygotes were used (*CX3CR1.Cre^ERT2^.hM4di.mCitrine* +/-) to minimize unwanted side effects as the Cre^ERT2^ transgene causes a functional null mutation of CX3CR1, an important regulator of microglial state, in the inserted allele. To confirm microglia-specific Gi DREADD expression, organotypic brain slice cultures (OBSCs) were prepared . Cre recombination was induced by addition of 4-OH-tamoxifen (TAM), which caused robust expression of the mCitrine tag within Iba-1+ microglia, indicated microglial-specific expression of the Gi DREADD (Figure 5A). Slices were then treated with a binge level concentration of ethanol (100mM) +/-the DREADD ligand CNO to activate Gi signaling in microglia. Gi DREADD activation in microglia blocked induction of proinflammatory genes by ethanol as reported^65^ (Figure 1D) as well as cell death (Figure 5B-C, F_3,19_=7.133, *p=*0.0021, \**p<*0.05, \*\**p<*0.01, Sidak’s post-test). We then assessed the role of microglia in ethanol-induced neurodegeneration *in vivo.* CNO was given 10 hours after ethanol *in vivo* to counter the induction of proinflammatory genes that occurs between 12 and 18 hours after binge ethanol^90^. Since the Gi DREADD mice are on the 129 background, rather than the B6J background used above, and these strains have different behavioral responses to ethanol^91^, we compared their blood ethanol concentrations (BECs) after 10 days of binge ethanol. C57BL/6J mice had an average BEC of 334 mg/dL 1 hour after ethanol, whereas the 129S mice had a BEC of 150.8 mg/dL. This mitigates concerns that the Gi DREADD mice might reach higher peak BECs. Ten days of binge ethanol caused robust TUNEL+ cell death in the OFC that was blocked by Gi DREADD inhibition of proinflammatory microglia (Figure 5D-E, F_3,16_=3.773, *p=*0.03, \**p<*0.05, Sidak’s post-test). This reveals a critical for role microglia in alcohol-induced neuronal death.

**Figure 5.**
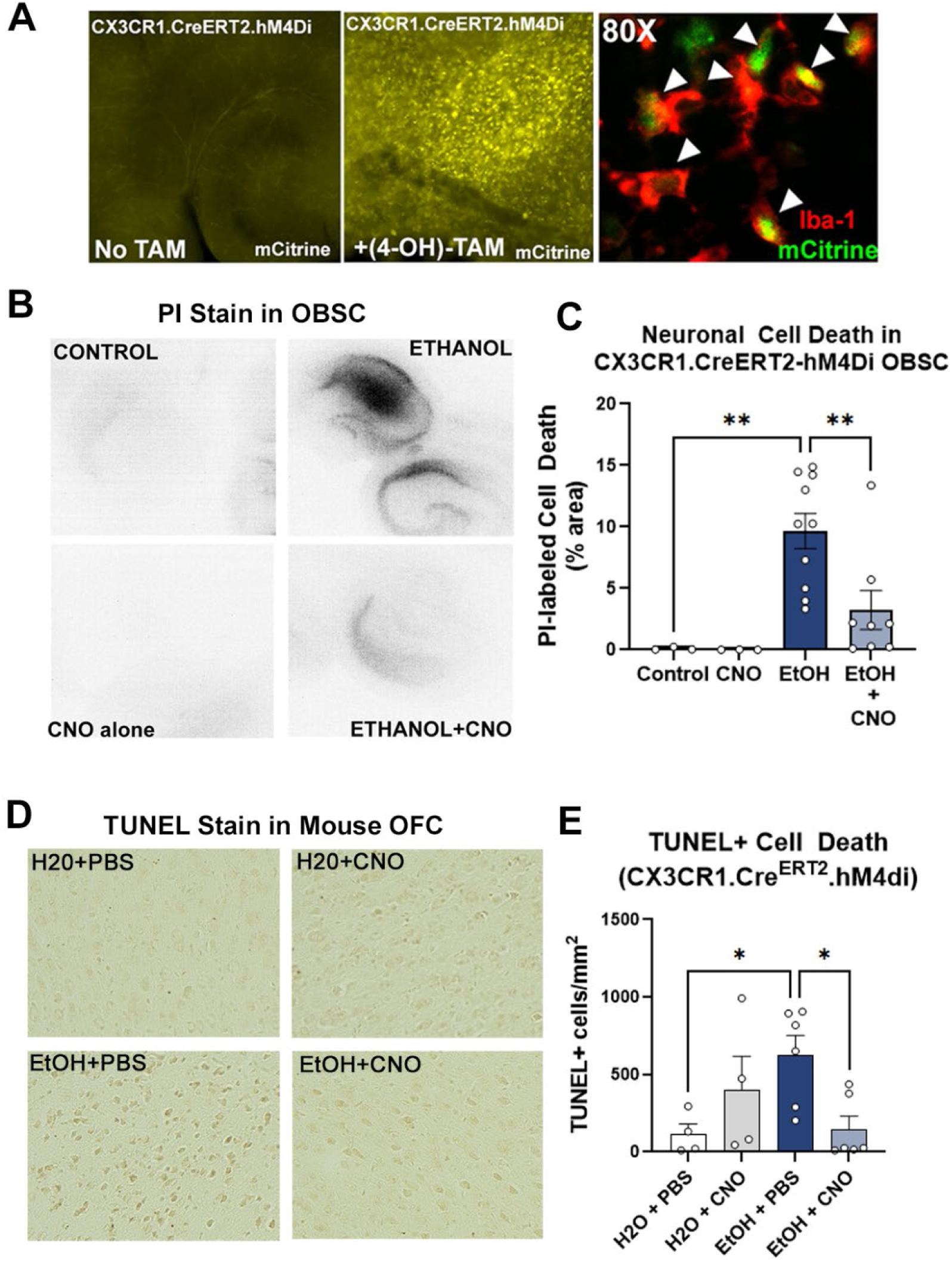
Gi DREADD inhibition of microglia blocks alcohol-induced neuronal cell death. (A-D) Organotypic brain slice cultures OBSCs from *CX3CR1.Cre^ERT2^.hM4di.mCitrine* (+/-) mice were used to assess the role of microglia on neuronal death *ex-vivo.* **(A)** Representative image +/-addition of 4-OH-TAM (10ng/mL) to induce Cre recombination which caused a robust increase in mCitrine within Iba-1+ microglia consistent with microglial DREADD expression. Arrows denote colocalization between Iba-1 and mCitrine. **(B)** Representative images of propidium iodide (PI) labeling of cell death in OBSC in response to ethanol. **(C)** Ethanol (100mM, 4 days) caused a robust increase in cell death 24 hours into withdrawal, which was significantly blocked by Gi signaling induced by CNO. 1-way ANOVA, F_3,19_=7.133, *p=*0.0021, \**p<*0.05, \*\**p<*0.01, Sidak’s post-test. N=3 Control slices, 3 CNO slices, 10 EtOH slices, 8 EtOH+CNO slices. **(D-E)** Induction of hM4Di Gi DREADD inhibition of microglia *in vivo*. *CX3CR1.Cre^ERT2^.hM4Di* (+/-) mice received 10 days of water or binge ethanol (5g/kg/d, i.g.) +/-CNO (3mg/kg, i.p., 10 hours after ethanol). Assessments were performed 24 hours following the final treatment. N= 4 H2O+PBS mice, 4 H2O+CNO mice, 6 EtOH+PBS mice, 6 EtOH+CNO mice. Males-open circles, females-filled circles. **(D)** Representative images of TUNEL cell death stain in OFC. **(E)** Ethanol increased TUNEL+ cell death by ∼3-fold , which was prevented by Gi DREADD inhibition of proinflammatory microglia. 1-way ANOVA, F_3,16_=3.773, *p=*0.03, \**p<*0.05, Sidak’s post-test.

### Inhibition of microglia prevents hyperkatifeia during abstinence in Gi DREADD mice

Given the persistence of proinflammatory microglial phenotypes and negative affective during prolonged withdrawal to binge ethanol, we assessed the impact of microglial inhibition the formation of hyperkatifeia phenotypes by ethanol. Female *CX3CR1.Cre^ERT2^.DIO.hM4di* (+/-) mice, showed similar to findings WT mice and protection by microglial inhibition. Anxiety-like and conditioned fear memory phenotypes were assessed 3 and 34 days into abstinence from 10 daily ethanol binges, respectively (Figure 6A). Consistent with findings above in male WT mice (Figure 3), ethanol increased anxiety-like behavior. Ethanol reduced center exploration 3 days into during abstinence with a trend toward a reduced distance traveled in the center of the open field (Figure 6B, F_2,20_=3.95, *p*=0.036, *p=*0.09) and reduced time spent in the center of the open field (Figure 6C, F_2,20_=7.1, *p*=0.0046, \**p<*0.05, Sidak’s post-test). Gi DREADD microglial inhibition in females prevented this measure of anxiety-like behavior, returning center distance traveled and time spent in the center of the open field to control levels (Figure 6B-C). Fine motor movement followed a similar pattern, with a trend toward a significant reduction after binge ethanol that returned to control levels with microglial inhibition (Figure 6D, F_2,21_=2.16, *p=*0.14). In conditioned fear memory testing, ethanol again had no impact on conditioned fear learning acquisition (Figure 6E). Ethanol modestly increased contextual fear memory during abstinence, with increased freezing during the last minute of assessment, which was prevented by microglial inhibition (Figure 6F). Similar to findings in male WT mice (Figure 3K), 10 ethanol binges in female Gi DREADD mice prevented the extinction of cued conditioned fear memory (Figure 6G). However, microglial inhibition during binge ethanol restored normal levels of fear memory extinction. Ten ethanol binges caused persistent increases in proinflammatory gene expression in female *CX3CR1.Cre^ERT2^.DIO.hM4di* (+/-) cortex (Figure 6H). Microglial Gi DREADD activation with CNO during binge ethanol significantly blunted much of this proinflammatory gene induction. Interestingly, time spent in the center was inversely correlated with *IFNα* (Figure 6I, Spearman’s R=-0.45). Thus, this approach to specifically inhibit proinflammatory microglial signaling during binge ethanol prevented persistent proinflammatory signaling and hyperkatifeia during abstinence in female Gi DREADD mice, implicating microglia in the development of these phenotypes.

**Figure 6.**
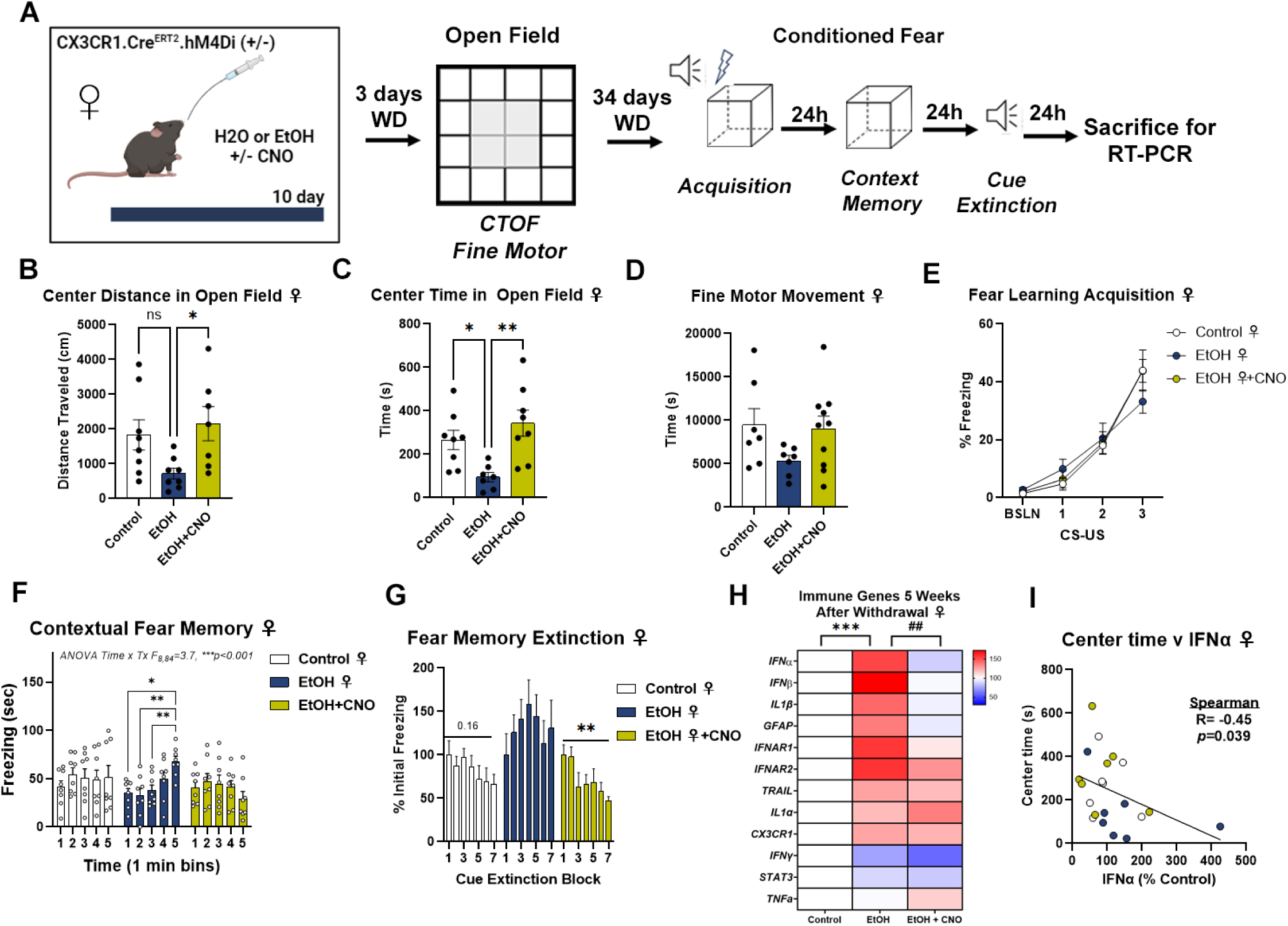
Gi DREADD inhibition of microglia attenuates alcohol-induced negative affect. **(A)** Experimental Design. Female *CX3CR1.Cre^ERT2^.hM4Di* (+/-) mice received 10 days of water or binge ethanol (5g/kg/d, i.g.) +/-CNO 10 hours after treatment. Mice underwent open field testing for anxiety-like behavior 3 days into withdrawal and conditioned fear for acquisition and extinction at 34 days into abstinence. Microglial inhibition prevented the induction of anxiety-like behavior caused by alcohol seen as a reduction in the (**B)** distance traveled in the center of the open field, 1-way ANOVA F_2,20_=3.95, *p*=0.036, ns-not significant, \**p<*0.05, Sidak’s post-test, and **(C)** reduced time spent in the center of the open field, 1-way ANOVA F_2,20_=7.1, *p*=0.0046, \*\**p<*0.01, \**p<*0.05, Sidak’s post-test. **(D)** Microglial inhibition prevented the trend toward a reduction in fine motor movement in the open field caused by ethanol. 1-was ANOVA F_2,21_=2.16, *p=*0.14. **(E)** Ethanol had no impact on conditioned fear learning acquisition. **(F)** In contextual fear memory assessments, ethanol increased freezing in the last minute of assessment that was prevented by microglial inhibition with CNO. 2-way ANOVA_time x treatment_ F_8,84_=3.7, *p<*0.0001, *\*p*<0.05, \*\**p<*0.01 Sidak’s post-test. **(G)** Ethanol disrupted the extinction of conditioned fear memory, which was restored by microglial inhibition. 1-way repeated measures ANOVAs: Control F_2.4,13.5_=2.1, *p=*0.16; EtOH F_2.8,19.8_=0.9, *p=*0.45; EtOH+CNO F_3.0,17.6_=6.6, \*\**p=*0.0035. **(H)** The persistent induction of multiple proinflammatory genes in the cortex 5 weeks into abstinence was prevented by Gi DREADD inhibition of microglia. 2-way ANOVA F_2,231_=6.921, *p=*0.001. \*\**p<*0.01, \**p<*0.05 vs control, #*#p<*0.01 vs EtOH. (I) A negative correlation between IFNα gene expression and Time spent in the center of the open field was found. IFN gene expression failed a normality test thus a Spearman correlation coefficient was calculated. R=-0.45, *p=*0.039. N= 8 female H2O+vehicle, 8 female EtOH+vehicle, and 8 female EtOH+CNO.

In male Gi DREADD mice (Supplemental Figure 2A), there were limitations to this approach including reduced tamoxifen-induced recombination. In males the hM4Di expression was <25% the level seen in females (Supplemental Figure 2B). Accordingly, though a trend toward a significant reduction in distance travelled in the center of the open field and time spent in the center was found in males, it was not impacted by microglial inhibition (Supplemental Figure 2C-D). Further, there was no impact of ethanol on fear learning acquisition (Supplemental Figure 2F), no changes was seen in contextual fear memory (Supplemental Figure 2F), and microglial inhibition had no effect on the extinction of cued conditioned memory in male mice receiving ethanol (Supplemental Figure 2H). Also, male Gi DREADD mice showed differences from WT mice during acute withdrawal as well as female Gi DREADD mice in that there were no signs of persistent proinflammatory signaling beyond an increase in GFAP (Supplemental Figure 2I).

## Discussion

AUD is a devastating disease with poor response rates to treatment and a high co-morbidity with affective disorders such as depression and PTSD^5,6,15–17^. Hyperkatifeia likely contributes to the maintenance of disease and relapse in AUD as well as to its psychiatric co-morbidities^7,8^. However, there are no available therapeutics that prevent or treat alcohol-induced hyperkatifeia. Thus, the present work on microglial mechanisms promoting hyperkatifeia is crucial to lay the groundwork for future studies elucidating therapeutic targets for this behavioral domain of AUD. Here, we found that 10-day binge ethanol induces neuronal death in the OFC, while causing immediate and persistent proinflammatory microglial polarization, loss of microglial BDNF in the dBNST, and long-lasting hyperkatifeia. Gi DREADD inhibition of proinflammatory microglia during repeated binge ethanol prevented OFC neuronal death and normalized the affective behavioral state, implicating microglia as key drivers of alcohol-induced neuroinflammation, neuronal death, and hyperkatifeia.

Prior reports have identified immediate increases in proinflammatory activation after binge ethanol, seen by induction of proinflammatory cytokines by way of the NF-kB pathway and increased gene expression^46,82,92,93^. However, few studies have assessed proinflammatory activation during prolonged abstinence in adults. The findings here of persistent proinflammatory activation weeks into abstinence are consistent with a recent report finding depressive behavior and increased cytokines 5 days into withdrawal from binge ethanol in PFC, hippocampus and striatum^94^, as well as other studies assessing brain cytokines at 3 and 5 days into withdrawal^79,95^. After adolescent binge ethanol, slight, yet persistent increases in proinflammatory cytokines are also found that are enhanced in Alzheimer’s disease models^69,96^. In humans that are abstinent for 18 days, plasma levels of IL-8, IL-1β, and IL-6 remained elevated, with IL-8 being correlated with craving^97^. However, this study finds a persistent induction of proinflammatory cytokines in cortex and microglial structural changes in stress-associated brain regions indicating long-lasting proinflammatory activation.

Microglia have been thought to be the main cell type responsible for proinflammatory signaling with alcohol abuse. This has been supported by studies using microglial depletion, which blunts much of the acute proinflammatory gene activation with binge ethanol^41,50^. Studies here further support a role for microglia in the initiation of proinflammatory signaling caused by binge alcohol. Using Gi DREADD microglia inhibition, which have been employed previously^57,58,65,89^, a reduction of persistent proinflammatory gene induction with ethanol was found. Particularly, type I IFN genes such as *IFNα*, and *IFNβ*, which were induced five weeks into withdrawal. *IFNα* was negatively correlated with center time (anxiety-like behavior) and fear extinction (PTSD-like behavioral inflexibility), suggesting IFN signaling might promote these behaviors. Indeed, IFN signaling promotes depressive phenotypes and extinction of conditioned fear memory in other settings^98–102^. Therefore, deciphering the contribution of IFN signaling to alcohol-induced negative affect is an important future direction. Together, these studies implicate a role for microglia in the persistent proinflammatory activation induced by ethanol, and suggest that persistent reprogramming of microglia to a proinflammatory phenotype contributes to the ongoing and persistent neurobiological dysfunction seen in AUD. Therefore, future studies should determine if normalization of persistent proinflammatory microglial phenotypes after binge ethanol can reverse the persistent negative affect during abstinence.

Microglial proinflammatory polarization has been known to be associated with alcohol-induced neuronal death^65,103–105^. This has been thought to contribute to alcohol-induced neurodegeneration^42,106,107^. However, a causative role for proinflammatory microglia in alcohol-induced neuronal death has only been recently uncovered. Carlson et al found that microglial depletion using colony stimulating factor receptor 1 (CSFR1) antagonism prevented alcohol-induced neuronal death caused by the Majchrowicz heavy continuous binge model^108^. The finding here that Gi DREADD inhibition of proinflammatory microglia blocks alcohol-induced neuronal death *ex-vivo* using brain slice cultures, and *in vivo* in the OFC, is consistent with that report. Further, it implicates a proinflammatory microglial phenotype in alcohol-induced neuronal death. Alcohol-induced neuronal death in the OFC could potentially contribute to the observed impairment in fear memory extinction, which is regulated in part by the OFC^54,109^. Future studies should directly test the role of OFC damage on behavioral flexibility in the setting of alcohol misuse. Previous reports found that alcohol promotes neuronal death through TRAIL-mediated apoptosis^47,63^. The current finding here of increased cleaved caspase-3 and TUNEL after alcohol is consistent with a potential role of apoptosis. Further, prior studies have suggested that microglia promote ethanol-induced neurodegeneration by the release of extracellular vesicles^45,110^. Thus, proinflammatory microglia may potentially promote neuronal death through their secretome.

The repeated binge ethanol treatment employed here models the alcohol pharmacokinetics seen in AUD. Cross-species pharmacokinetic comparisons find the 5g/kg gavage equates to ∼5-7 drinks in adult humans, which are easily achieved by individuals with AUD^60^. The proinflammatory microglial polarization and neuronal death observed in the OFC are also found in human AUD brain^47,82^ supporting translational validity of the model. However, there are some limitations to non-contingent ethanol administration paradigms such as the potential for gavage stress. Proinflammatory gene induction and microglial proinflammatory signaling has also been observed in alcohol self-administration models such as drinking in the dark (DID) which achieve lower blood alcohol levels (typical BACs<0.1mg/dL)^111–113^. Neuronal death was not seen in the amygdala after 3 cycles of DID using Flourojade B, however, the OFC was not assessed^111^. Though no studies to date have performed a comparable 10 day DID study, it is likely that the repeated exposure to higher blood ethanol concentrations that are achieved in AUD, and the 5g/kg gavage model are required for neuronal death. This is supported by *ex-vivo* studies that find neuronal death with comparable alcohol concentrations yet lack other potential contributors such as gavage stress^47^. Gavage stress is an important consideration, however, particularly with the first ethanol exposure. Findings from the 4-day gavage binge ethanol group, which showed no hyperkatifeia-associated behavioral differences, argue that gavage stress does not drive the long-lasting behavioral changes seen in this model. Rather, it appears that repeated binge ethanol exposures drive these effects, as has been postulated previously^114^. However, it would be of interest for future studies to determine the threshold of alcohol exposure, needed *in vivo* to result in neuronal death and behavioral dysfunction, either with contingent or non-contingent administration. Since this study now provides direct evidence that proinflammatory microglia promote ethanol-induced neurodegeneration *in vivo* and *ex-vivo*, as well as persistent hyperkatifeia, future work should further define these microglia-specific mechanisms to identify druggable microglia-specific therapeutic targets.

The prevention of negative affective/hyperkatifeia phenotypes by the inhibition of proinflammatory microglia in female Gi DREADD mice strongly implicates their role in these persistent maladaptive behavioral phenotypes. Findings of reduced fear memory extinction with alcohol are consistent with a report that homeostatic microglia assist in the forgetting of conditioned fear memory^37^. On the other hand, prior reports identified protective features of regenerative microglia, which repopulate the brain after CSFR1-mediated microglial depletion^115,116^. Regenerative microglia have a more trophic phenotype (e.g., increased BDNF) and show reduced proinflammatory responses to alcohol^57,65^. These findings suggest that proinflammatory microglia may lose important trophic (e.g., BDNF) or other neuronal modulatory functions that promote fear memory extinction through modulation of neuronal plasticity. BDNF is altered in AUD and is related to AUD behavioral phenotypes. Previous studies have linked excessive drinking in rats with lower levels of BDNF in the BNST and amygdala^117,118^. Additional work found that knocking down BDNF in the CeA with antisense oligonucleotides increases anxiety-like behavior and alcohol self-administration^34^. Further, in human plasma BDNF is associated with functional connectivity between the amygdala and prefrontal cortex^118^. The finding here of reduced microglial BDNF in the BNST 5 weeks into abstinence raises the question as to whether BDNF regulates neuronal activity in this region. Indeed, a recent study found that BDNF application reduces excitability of neurons in the BNST, with BDNF/TrkB signaling needed for LTD in BNST neurons^119^. Considering the role of BDNF in neuronal function and negative affect, future studies should examine if microglial BDNF loss promotes hyperkatifeia.

The findings that chemogenetic microglial inhibition prevented anxiety-like and fear memory phenotypes are consistent with work using microglial depletion, which prevented the alcohol dependence-induced escalation of drinking and reduced sucrose intake, a measure of anhedonia^51^. However, it is not clear whether inhibiting microglia after binge alcohol-induced negative affect or dependence-induced escalation of drinking would reverse these AUD phenotypes. These are important future directions. The inability of microglial inhibition to improve affective responses in male Gi DREADD mice is confounded by the low level of hM4di expression. Thus, it is difficult to know if this represents a true sex difference in the role of microglia, or if the approach had inherent limitations in males. Both male WT and female *CX3CR1.Cre^ERT2^.hM4di* showed similar OFC neuronal death, proinflammatory gene induction, anxiety-like behavior, and reduced fear memory extinction with 10 days of binge ethanol. This suggests that limitations of this approach in the male Gi DREADD mice may underlie the lack of protection of microglial inhibition. However, this remains unclear given the current confounding factors. Given there may be differences in sensitivity to the effects of alcohol between the sexes, comprehensive studies investigating the sexually dimorphic immune responses across multiple brain regions and strains would be of value to the field. However, the data here find that with severe features such as neuronal death and persistent negative affect that both sexes are similarly impacted. Further, it is important to note that the approach using the CX3CR1 promoter had some key limitations that also relate to sex. First, the *DIO.hM4di* mice are on the 129 background, which has different responses to ethanol than B6J mice^91^. Also, in this model, a copy of CX3CR1, a G-coupled protein receptor with anti-inflammatory properties^120^, is lost with Cre gene insertion. Therefore, an alternative microglial specific promoter such as Tmem119, is likely a better approach for microglial inhibition to effectively elucidate mechanisms driving ethanol-induced deficits in complex behaviors.

There are additional important limitations to this study. First, proinflammatory gene expression was measured in whole frontal cortex. Assessment of individual frontal cortical regions (e.g., prelimbic or infralimbic) as well as subcortical regions (e.g., dBNST or CeA) using tissue punches would be of value. Though we report center time and center distance, the number of center entries in the open field is a useful measure that was not captured by our software. Further, the chemogenetic approach to inhibit microglia is limited in that it impacts the entire brain. There are significant challenges for brain regional inhibition of microglia given their inherent difficulty to transfect with viruses. However, as more viruses emerge that are capable of generating specific expression in microglia without inducing robust immune responses, it will become more feasible to regulate microglia in specific brain regions.

In summary, we find that chronic binge ethanol induces proinflammatory microglial polarization and persistent loss of microglial BDNF in hyperkatifeia-related brain regions, OFC neuronal cell death, and negative affect during acute withdrawal and prolonged abstinence. These neuropathological features were blocked by chemogenetic inhibition of microglia, implicating microglia as drivers of alcohol-induced hyperkatifeia. The translational value of this work is that it implicates microglia in key AUD-associated pathology and warrants investigation of microglia-specific therapeutic targets. Future studies will define the specific molecular mechanisms by which microglia promote these negative-affect states and other related AUD phenotypes to identify novel therapeutic targets.

## Acknowledgements

We thank Jay Campbell for his help with laboratory management. Paige Anton assisted with editing the manuscript. The senior author LGC thanks God the Father for his grace and guidance.

## Funding sources

This work was supported by the National Institutes of Health AA024829 (LGC), AA028924 (LGC), AA031414 (LGC), AA011605 (LGC), AA030463 (LGC), P50HD103573 (SM), and the Bowles Center for Alcohol Studies.

## Declarations of Interest

None

---

**Supplemental Figure 1:**
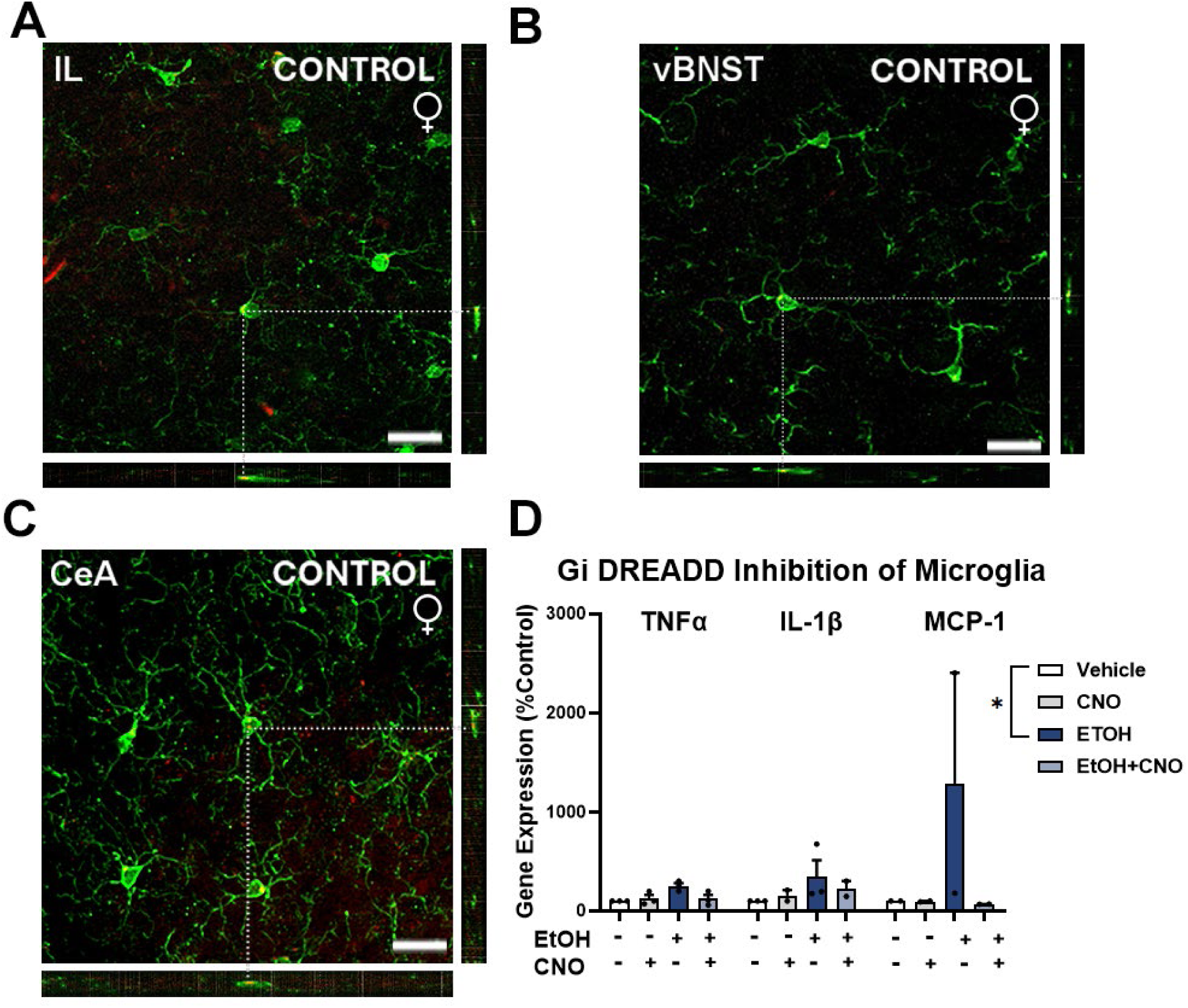
**(A-D)** Orthogonal projections of confocal images for Iba-1 and BDNF co-IF twenty-four hours after 10 ethanol binges confirming colocalization in the **(A)** IL, **(B)** vBNST, and **(C)** CeA. **(D)** Organotypic brain slice cultures OBSCs from CX3CR1.Cre^ERT2^.hM4di.mCitrine (+/-) mice were used to assess the role of microglia on proinflammatory gene expression and neuronal death *ex-vivo.* Ethanol (100µM, 96 hours) caused increases in proinflammatory gene expression (TNFα, IL-1β, and MCP-1) which was blocked by Gi DREADD signaling induced by CNO (5µM). 2-way ANOVA, Treatment effect: F_3,18_=3.14, *p=*0.044. *#p<*0.05, Dunnett’s multiple comparisons. Each data point represents the average of 2 technical replicates for 2-3 experimental replicates (i.e., separate slice culture preparations) for each gene assessed.

**Supplemental Figure 2.**
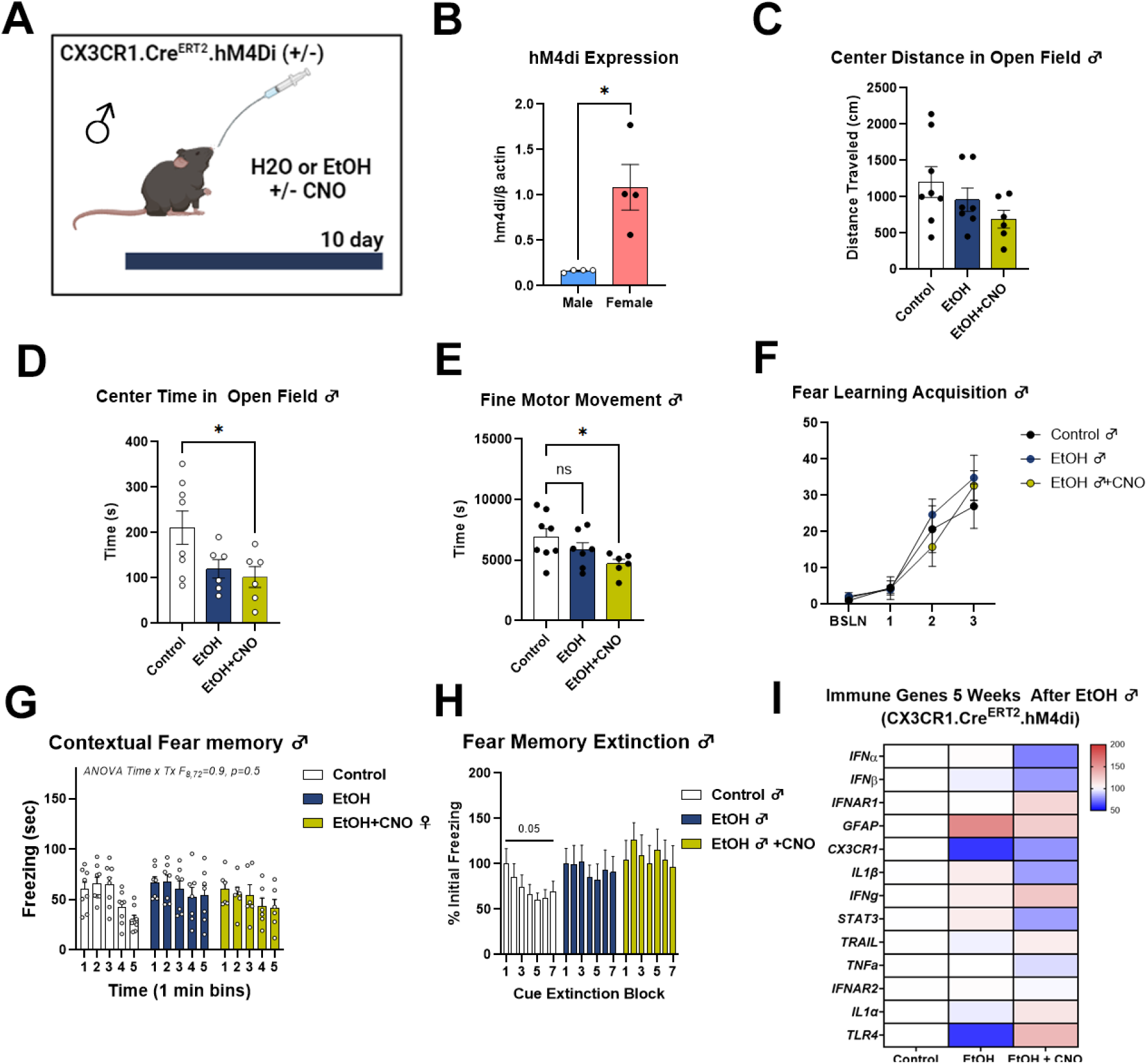
Gi DREADD inhibition of microglia fails to attenuate alcohol-induced negative affect in males. **(A)** Experimental Design. Male *CX3CR1.Cre^ERT2^.hM4Di* (+/-) mice received 10 days of water or binge ethanol (5g/kg/d, i.g.) +/-CNO 10 hours after treatment. Mice underwent open field testing for anxiety-like behavior 3 days into withdrawal and conditioned fear for acquisition and extinction at 34 days into abstinence. **(B)** hM4di expression was measured by RT-PCR. Males showed ∼25% of the level hM4di expression as females. Microglial inhibition failed to prevent the induction of anxiety-like behavior caused by alcohol seen change in the (**C)** distance traveled in the center of the open field, 1-way ANOVA F_2,18_=2.0, *p*=0.16 and **(D)** a reduction in the time spent in the center of the open field, 1-way ANOVA F_2,17_=3.99, *p*=0.038, \**p<*0.05, Sidak’s post-test. **(E)** Fine motor movement was significantly reduced in the EtOH+CNO group. 1-way ANOVA F_2,18_=3.5, *p=*0.05, ns-not significant, \**p<*0.05, Sidak’s post-test. **(F)** Ethanol had no impact on conditioned fear memory acquisition.. **(G)** Neither ethanol nor microglial inhibition had any effect on contextual fear memory. 2-way ANOVA_time x treatment_ F_8,72_=0.9, *p=*0.5. **(H)** Microglial inhibition had no impact on fear memory extinction. 1-way repeated measures ANOVAs: Control F_2.86,16.19_=3.2, *p=*0.053; EtOH F_2.999,17.99_=0.38, *p=*0.77; EtOH+CNO F_1.94,11.7_=0.9, *p=*0.43. **(I)** No robust proinflammatory gene induction signature was seen with EtOH in the hM4di mice, with little impact of microglial inhibition.

## Notes

### Competing Interest Statement

The authors have declared no competing interest.

### Summary of Updates

This manuscript has been revised to increase the N for sex comparisons in microglia structure across brain regions (revised Figure 2). Neuronal activation data has been removed.

## References

1 Whiteford, H. A., Ferrari, A. J., Degenhardt, L., Feigin, V. & Vos, T. The global burden of mental, neurological and substance use disorders: an analysis from the Global Burden of Disease Study 2010. PloS one 10, e0116820 (2015). 10.1371/journal.pone.0116820

2 Goldstein, R. B. et al. Nosologic Comparisons of DSM-IV and DSM-5 Alcohol and Drug Use Disorders: Results From the National Epidemiologic Survey on Alcohol and Related Conditions-III. J Stud Alcohol Drugs 76, 378–388 (2015). 10.15288/jsad.2015.76.378

3 Burnette, E. M. et al. Novel Agents for the Pharmacological Treatment of Alcohol Use Disorder. Drugs 82, 251–274 (2022). 10.1007/s40265-021-01670-3

4 Anderson, P., O’Donnell, A. & Kaner, E. Managing Alcohol Use Disorder in Primary Health Care. Curr Psychiatry Rep 19, 79 (2017). 10.1007/s11920-017-0837-z

5 Whitford, J. L., Widner, S. C., Mellick, D. & Elkins, R. L. Self-report of drinking compared to objective markers of alcohol consumption. Am J Drug Alcohol Abuse 35, 55–58 (2009). 10.1080/00952990802295212

6 Maisto, S. A., Pollock, N. K., Cornelius, J. R., Lynch, K. G. & Martin, C. S. Alcohol relapse as a function of relapse definition in a clinical sample of adolescents. Addict Behav 28, 449–459 (2003). 10.1016/s0306-4603(01)00267-2

7 Koob, G. F. Anhedonia, Hyperkatifeia, and Negative Reinforcement in Substance Use Disorders. Curr Top Behav Neurosci 58, 147–165 (2022). 10.1007/7854_2021_288

8 Koob, G. F. Drug Addiction: Hyperkatifeia/Negative Reinforcement as a Framework for Medications Development. Pharmacological reviews 73, 163–201 (2021). 10.1124/pharmrev.120.000083

9 Sliedrecht, W., de Waart, R., Witkiewitz, K. & Roozen, H. G. Alcohol use disorder relapse factors: A systematic review. Psychiatry Res 278, 97–115 (2019). 10.1016/j.psychres.2019.05.038

10 Sinha, R. How does stress increase risk of drug abuse and relapse? Psychopharmacology 158, 343–359 (2001). 10.1007/s002130100917

11 Greenfield, S. F. et al. The effect of depression on return to drinking: a prospective study. Archives of general psychiatry 55, 259–265 (1998). 10.1001/archpsyc.55.3.259

12 Cooper, M. L., Russell, M., Skinner, J. B., Frone, M. R. & Mudar, P. Stress and alcohol use: moderating effects of gender, coping, and alcohol expectancies. J Abnorm Psychol 101, 139–152 (1992). 10.1037//0021-843x.101.1.139

13 Brown, S. A., Vik, P. W., Patterson, T. L., Grant, I. & Schuckit, M. A. Stress, vulnerability and adult alcohol relapse. J Stud Alcohol 56, 538–545 (1995). 10.15288/jsa.1995.56.538

14 Sinha, R. et al. Enhanced negative emotion and alcohol craving, and altered physiological responses following stress and cue exposure in alcohol dependent individuals. Neuropsychopharmacology : official publication of the American College of Neuropsychopharmacology 34, 1198–1208 (2009). 10.1038/npp.2008.78

15 Blanco, C. et al. Comorbidity of posttraumatic stress disorder with alcohol dependence among US adults: results from National Epidemiological Survey on Alcohol and Related Conditions. Drug and alcohol dependence 132, 630–638 (2013). 10.1016/j.drugalcdep.2013.04.016

16 Gielen, N., Havermans, R. C., Tekelenburg, M. & Jansen, A. Prevalence of post-traumatic stress disorder among patients with substance use disorder: it is higher than clinicians think it is. Eur J Psychotraumatol 3 (2012). 10.3402/ejpt.v3i0.17734

17 Read, J. P., Brown, P. J. & Kahler, C. W. Substance use and posttraumatic stress disorders: symptom interplay and effects on outcome. Addict Behav 29, 1665–1672 (2004). 10.1016/j.addbeh.2004.02.061

18 Keyes, K. M., Hatzenbuehler, M. L. & Hasin, D. S. Stressful life experiences, alcohol consumption, and alcohol use disorders: the epidemiologic evidence for four main types of stressors. Psychopharmacology (Berl) 218, 1–17 (2011). 10.1007/s00213-011-2236-1

19 Fetzner, M. G., McMillan, K. A., Sareen, J. & Asmundson, G. J. What is the association between traumatic life events and alcohol abuse/dependence in people with and without PTSD? Findings from a nationally representative sample. Depress Anxiety 28, 632–638 (2011). 10.1002/da.20852

20 Huang, M. C., Schwandt, M. L., Ramchandani, V. A., George, D. T. & Heilig, M. Impact of multiple types of childhood trauma exposure on risk of psychiatric comorbidity among alcoholic inpatients. Alcoholism, clinical and experimental research 36, 1099–1107 (2012). 10.1111/j.1530-0277.2011.01695.x

21 Elsayed, N. M. et al. Trajectories of Alcohol Initiation and Use During Adolescence: The Role of Stress and Amygdala Reactivity. J Am Acad Child Adolesc Psychiatry 57, 550–560 (2018). 10.1016/j.jaac.2018.05.011

22 Lê, A. D. et al. Reinstatement of alcohol-seeking by priming injections of alcohol and exposure to stress in rats. Psychopharmacology 135, 169–174 (1998). 10.1007/s002130050498

23 Breese, G. R., Overstreet, D. H. & Knapp, D. J. Conceptual framework for the etiology of alcoholism: a “kindling”/stress hypothesis. Psychopharmacology (Berl) 178, 367–380 (2005). 10.1007/s00213-004-2016-2

24 Marcinkiewcz, C. A., Dorrier, C. E., Lopez, A. J. & Kash, T. L. Ethanol induced adaptations in 5-HT2c receptor signaling in the bed nucleus of the stria terminalis: implications for anxiety during ethanol withdrawal. Neuropharmacology 89, 157–167 (2015). 10.1016/j.neuropharm.2014.09.003

25 Delfs, J. M., Zhu, Y., Druhan, J. P. & Aston-Jones, G. Noradrenaline in the ventral forebrain is critical for opiate withdrawal-induced aversion. Nature 403, 430–434 (2000). 10.1038/35000212

26 Pati, D. et al. Chronic intermittent ethanol exposure dysregulates a GABAergic microcircuit in the bed nucleus of the stria terminalis. Neuropharmacology 168, 107759 (2020). 10.1016/j.neuropharm.2019.107759

27 Sink, K. S. et al. Effects of continuously enhanced corticotropin releasing factor expression within the bed nucleus of the stria terminalis on conditioned and unconditioned anxiety. Mol Psychiatry 18, 308–319 (2013). 10.1038/mp.2011.188

28 Walker, D. L., Toufexis, D. J. & Davis, M. Role of the bed nucleus of the stria terminalis versus the amygdala in fear, stress, and anxiety. European journal of pharmacology 463, 199–216 (2003). 10.1016/s0014-2999(03)01282-2

29 LaBar, K. S., Gatenby, J. C., Gore, J. C., LeDoux, J. E. & Phelps, E. A. Human amygdala activation during conditioned fear acquisition and extinction: a mixed-trial fMRI study. Neuron 20, 937–945 (1998). 10.1016/s0896-6273(00)80475-4

30 Flores-Ramirez, F. J. et al. Blockade of orexin receptors in the infralimbic cortex prevents stress-induced reinstatement of alcohol-seeking behaviour in alcohol-dependent rats. British journal of pharmacology 180, 1500–1515 (2023). 10.1111/bph.16015

31 Flores-Ramirez, F. J., Matzeu, A., Sánchez-Marín, L. & Martin-Fardon, R. Blockade of corticotropin-releasing factor-1 receptors in the infralimbic cortex prevents stress-induced reinstatement of alcohol seeking in male Wistar rats: Evidence of interaction between CRF. Neuropharmacology 210, 109046 (2022). 10.1016/j.neuropharm.2022.109046

32 Breese, G. R. et al. Repeated lipopolysaccharide (LPS) or cytokine treatments sensitize ethanol withdrawal-induced anxiety-like behavior. Neuropsychopharmacology 33, 867–876 (2008). 10.1038/sj.npp.1301468

33 Duivis, H. E., Vogelzangs, N., Kupper, N., de Jonge, P. & Penninx, B. W. Differential association of somatic and cognitive symptoms of depression and anxiety with inflammation: findings from the Netherlands Study of Depression and Anxiety (NESDA). Psychoneuroendocrinology 38, 1573–1585 (2013). 10.1016/j.psyneuen.2013.01.002

34 Pandey, S. C., Zhang, H., Roy, A. & Misra, K. Central and medial amygdaloid brain-derived neurotrophic factor signaling plays a critical role in alcohol-drinking and anxiety-like behaviors. The Journal of neuroscience : the official journal of the Society for Neuroscience 26, 8320–8331 (2006). 10.1523/jneurosci.4988-05.2006

35 Silva-Peña, D. et al. Alcohol-induced cognitive deficits are associated with decreased circulating levels of the neurotrophin BDNF in humans and rats. Addict Biol 24, 1019–1033 (2019). 10.1111/adb.12668

36 Yu, Y., Yuan, Z., Fan, Y., Li, J. & Wu, Y. Dynamic Transitions in Neuronal Network Firing Sustained by Abnormal Astrocyte Feedback. Neural plasticity 2020, 8864246 (2020). 10.1155/2020/8864246

37 Wang, C. et al. Microglia mediate forgetting via complement-dependent synaptic elimination. Science 367, 688–694 (2020). 10.1126/science.aaz2288

38 Lan, L. et al. Chronic exposure of alcohol triggers microglia-mediated synaptic elimination inducing cognitive impairment. Experimental neurology 353, 114061 (2022). 10.1016/j.expneurol.2022.114061

39 Peng, H., Geil Nickell, C. R., Chen, K. Y., McClain, J. A. & Nixon, K. Increased expression of M1 and M2 phenotypic markers in isolated microglia after four-day binge alcohol exposure in male rats. Alcohol 62, 29–40 (2017). 10.1016/j.alcohol.2017.02.175

40 Crews, F. T., Zou, J. & Coleman, L. G., Jr. Extracellular microvesicles promote microglia-mediated pro-inflammatory responses to ethanol. Journal of neuroscience research 99, 1940–1956 (2021). 10.1002/jnr.24813

41 Walter, T. J. & Crews, F. T. Microglial depletion alters the brain neuroimmune response to acute binge ethanol withdrawal. Journal of neuroinflammation 14, 86 (2017). 10.1186/s12974-017-0856-z

42 Zou, J. & Crews, F. Induction of innate immune gene expression cascades in brain slice cultures by ethanol: key role of NF-κB and proinflammatory cytokines. Alcoholism, clinical and experimental research 34, 777–789 (2010). 10.1111/j.1530-0277.2010.01150.x

43 Crews, F. T., Lawrimore, C. J., Walter, T. J. & Coleman, L. G., Jr. The role of neuroimmune signaling in alcoholism. Neuropharmacology 122, 56–73 (2017). 10.1016/j.neuropharm.2017.01.031

44 Vetreno, R. P., Qin, L., Coleman, L. G. & Crews, F. T. Increased Toll-like Receptor-MyD88-NFκB-Proinflammatory neuroimmune signaling in the orbitofrontal cortex of humans with alcohol use disorder. Alcohol Clin Exp Res 45, 1747–1761 (2021). 10.1111/acer.14669

45 Coleman, L. G., Zou, J. & Crews, F. T. Microglial-derived miRNA let-7 and HMGB1 contribute to ethanol-induced neurotoxicity via TLR7. J Neuroinflammation 14, 22 (2017). 10.1186/s12974-017-0799-4

46 Crews, F. T., Qin, L., Sheedy, D., Vetreno, R. P. & Zou, J. High mobility group box 1/Toll-like receptor danger signaling increases brain neuroimmune activation in alcohol dependence. Biological psychiatry 73, 602–612 (2013). 10.1016/j.biopsych.2012.09.030

47 Qin, L. et al. TRAIL Mediates Neuronal Death in AUD: A Link between Neuroinflammation and Neurodegeneration. International journal of molecular sciences 22 (2021). 10.3390/ijms22052547

48 Lovelock, D. F. et al. The Toll-like receptor 7 agonist imiquimod increases ethanol self-administration and induces expression of Toll-like receptor related genes. Addict Biol 27, e13176 (2022). 10.1111/adb.13176

49 Lovelock, D. F. et al. Increased alcohol self-administration following repeated Toll-like receptor 3 agonist treatment in male and female rats. Pharmacol Biochem Behav 216, 173379 (2022). 10.1016/j.pbb.2022.173379

50 Warden, A. S. et al. Microglia Control Escalation of Drinking in Alcohol-Dependent Mice: Genomic and Synaptic Drivers. Biol Psychiatry 88, 910–921 (2020). 10.1016/j.biopsych.2020.05.011

51 Warden, A. S. et al. Microglia depletion and alcohol: Transcriptome and behavioral profiles. Addict Biol 26, e12889 (2021). 10.1111/adb.12889

52 Koob, G. F. & Volkow, N. D. Neurobiology of addiction: a neurocircuitry analysis. The lancet. Psychiatry 3, 760–773 (2016). 10.1016/S2215-0366(16)00104-8

53 Zois, E. et al. Orbitofrontal structural markers of negative affect in alcohol dependence and their associations with heavy relapse-risk at 6 months post-treatment. European psychiatry : the journal of the Association of European Psychiatrists 46, 16–22 (2017). 10.1016/j.eurpsy.2017.07.013

54 Hamilton, D. A. & Brigman, J. L. Behavioral flexibility in rats and mice: contributions of distinct frontocortical regions. Genes Brain Behav 14, 4–21 (2015). 10.1111/gbb.12191

55 VanElzakker, M. B., Dahlgren, M. K., Davis, F. C., Dubois, S. & Shin, L. M. From Pavlov to PTSD: the extinction of conditioned fear in rodents, humans, and anxiety disorders. Neurobiology of learning and memory 113, 3–18 (2014). 10.1016/j.nlm.2013.11.014

56 Zou, J. et al. Ethanol Induces Secretion of Proinflammatory Extracellular Vesicles That Inhibit Adult Hippocampal Neurogenesis Through G9a/GLP-Epigenetic Signaling. Front Immunol 13, 866073 (2022). 10.3389/fimmu.2022.866073

57 Zou, J. et al. Microglia either promote or restrain TRAIL-mediated excitotoxicity caused by Aβ. J Neuroinflammation 21, 215 (2024). 10.1186/s12974-024-03208-2

58 Grace, P. M. et al. DREADDed microglia in pain: Implications for spinal inflammatory signaling in male rats. Experimental neurology 304, 125–131 (2018). 10.1016/j.expneurol.2018.03.005

59 Bertola, A., Mathews, S., Ki, S. H., Wang, H. & Gao, B. Mouse model of chronic and binge ethanol feeding (the NIAAA model). Nat Protoc 8, 627–637 (2013). 10.1038/nprot.2013.032

60 Pruett, S., Tan, W., Howell, G. E., 3rd & Nanduri, B. Dosage scaling of alcohol in binge exposure models in mice: An empirical assessment of the relationship between dose, alcohol exposure, and peak blood concentrations in humans and mice. Alcohol 89, 9–17 (2020). 10.1016/j.alcohol.2020.03.011

61 Barnett, A. M. et al. Loss of neuronal lysosomal acid lipase drives amyloid pathology in Alzheimer’s disease. bioRxiv (2024). 10.1101/2024.06.09.596693

62 Stoppini, L., Buchs, P. A. & Muller, D. A simple method for organotypic cultures of nervous tissue. J Neurosci Methods 37, 173–182 (1991).

63 Zou, J. et al. Microglia either promote or restrain TRAIL-mediated excitotoxicity caused by Abeta(1-42) oligomers. Journal of neuroinflammation 21, 215 (2024). 10.1186/s12974-024-03208-2

64 Zou, J. et al. Ethanol Induces Secretion of Proinflammatory Extracellular Vesicles That Inhibit Adult Hippocampal Neurogenesis Through G9a/GLP-Epigenetic Signaling. Frontiers in immunology 13 (2022). 10.3389/fimmu.2022.866073

65 Coleman, L. G., Jr., Zou, J. & Crews, F. T. Microglial depletion and repopulation in brain slice culture normalizes sensitized proinflammatory signaling. Journal of neuroinflammation 17, 27 (2020). 10.1186/s12974-019-1678-y

66 Zou, J. Y. & Crews, F. T. Release of neuronal HMGB1 by ethanol through decreased HDAC activity activates brain neuroimmune signaling. PloS one 9, e87915 (2014). 10.1371/journal.pone.0087915

67 Zou, J. & Crews, F. CREB and NF-kappaB transcription factors regulate sensitivity to excitotoxic and oxidative stress induced neuronal cell death. Cellular and molecular neurobiology 26, 385–405 (2006). 10.1007/s10571-006-9045-9

68 Zou, J. Y. & Crews, F. T. TNF alpha potentiates glutamate neurotoxicity by inhibiting glutamate uptake in organotypic brain slice cultures: neuroprotection by NF kappa B inhibition. Brain research 1034, 11–24 (2005). 10.1016/j.brainres.2004.11.014

69 Barnett, A. et al. Adolescent Binge Alcohol Enhances Early Alzheimer’s Disease Pathology in Adulthood Through Proinflammatory Neuroimmune Activation. Front Pharmacol 13, 884170 (2022). 10.3389/fphar.2022.884170

70 Wang, X., Spandidos, A., Wang, H. & Seed, B. PrimerBank: a PCR primer database for quantitative gene expression analysis, 2012 update. Nucleic acids research 40, D1144–1149 (2012). 10.1093/nar/gkr1013

71 Wang, W. et al. Microglial repopulation reverses cognitive and synaptic deficits in an Alzheimer’s disease model by restoring BDNF signaling. Brain, behavior, and immunity 113, 275–288 (2023). 10.1016/j.bbi.2023.07.011

72 McIlwain, K. L., Merriweather, M. Y., Yuva-Paylor, L. A. & Paylor, R. The use of behavioral test batteries: effects of training history. Physiol Behav 73, 705–717 (2001). 10.1016/s0031-9384(01)00528-5

73. Paylor, R., Spencer, C. M., Yuva-Paylor, L. A. & Pieke-Dahl, S. The use of behavioral test batteries, II: effect of test interval. *Physiol Behav* 87, 95-102 (2006). 10.1016/j.physbeh.2005.09.002

74 Barnett, A. G. et al. Loss of neuronal lysosomal acid lipase contributes to Alzheimer’s disease pathology and cognitive decline. Alzheimer’s & dementia : the journal of the Alzheimer’s Association (2025). 10.1002/alz.70486

75 Barnett, A. M. et al. Adolescent Binge Alcohol Enhances Early Alzheimer’s Disease Pathology in Adulthood Through Proinflammatory Neuroimmune Activation. Frontiers in pharmacology 13, 884170 (2022). 10.3389/fphar.2022.884170

76 Fish, E. W. et al. The enduring impact of neurulation stage alcohol exposure: A combined behavioral and structural neuroimaging study in adult male and female C57BL/6J mice. Behavioural brain research 338, 173–184 (2018). 10.1016/j.bbr.2017.10.020

77 Coleman, L. G., Jr., Crews, F. T. & Vetreno, R. P. The persistent impact of adolescent binge alcohol on adult brain structural, cellular, and behavioral pathology: A role for the neuroimmune system and epigenetics. International review of neurobiology 160, 1–44 (2021). 10.1016/bs.irn.2021.08.001

78 Crews, F., Qin, L., Coleman Jr, L., Vidrascu, E. & Vetreno, R. Cortical reactive microglia activate astrocytes, increasing neurodegeneration in human alcohol use disorder. BioRxiv (2025). 10.1101/2025.04.25.650687

79 Qin, L. & Crews, F. T. Chronic ethanol increases systemic TLR3 agonist-induced neuroinflammation and neurodegeneration. Journal of neuroinflammation 9, 130 (2012). 10.1186/1742-2094-9-130

80 Qin, L. et al. Increased systemic and brain cytokine production and neuroinflammation by endotoxin following ethanol treatment. Journal of neuroinflammation 5, 10 (2008). 10.1186/1742-2094-5-10

81 Miguel-Hidalgo, J. J., Overholser, J. C., Meltzer, H. Y., Stockmeier, C. A. & Rajkowska, G. Reduced glial and neuronal packing density in the orbitofrontal cortex in alcohol dependence and its relationship with suicide and duration of alcohol dependence. Alcoholism, clinical and experimental research 30, 1845–1855 (2006). 10.1111/j.1530-0277.2006.00221.x

82 He, J. & Crews, F. T. Increased MCP-1 and microglia in various regions of the human alcoholic brain. Exp Neurol 210, 349–358 (2008). 10.1016/j.expneurol.2007.11.017

83 Melbourne, J. K., Wooden, J. I., Carlson, E. R., Anasooya Shaji, C. & Nixon, K. Neuroimmune Activation and Microglia Reactivity in Female Rats Following Alcohol Dependence. International journal of molecular sciences 25 (2024). 10.3390/ijms25031603

84 Godeanu, S. & Catalin, B. The Complementary Role of Morphology in Understanding Microglial Functional Heterogeneity. International journal of molecular sciences 26 (2025). 10.3390/ijms26083811

85 (!!! INVALID CITATION !!! 26–30).

86 Crews, F. T. & Vetreno, R. P. Mechanisms of neuroimmune gene induction in alcoholism. Psychopharmacology 233, 1543–1557 (2016). 10.1007/s00213-015-3906-1

87 Kettenmann, H., Hanisch, U. K., Noda, M. & Verkhratsky, A. Physiology of microglia. Physiol Rev 91, 461–553 (2011). 10.1152/physrev.00011.2010

88 Pan, K. & Garaschuk, O. The role of intracellular calcium-store-mediated calcium signals in in vivo sensor and effector functions of microglia. J Physiol 601, 4203–4215 (2023). 10.1113/JP279521

89 Parusel, S., Yi, M. H., Hunt, C. L. & Wu, L. J. Chemogenetic and Optogenetic Manipulations of Microglia in Chronic Pain. Neuroscience bulletin 39, 368–378 (2023). 10.1007/s12264-022-00937-3

90 Walter, T. J. & Crews, F. T. Microglial depletion alters the brain neuroimmune response to acute binge ethanol withdrawal. Journal of neuroinflammation 14, 86 (2017). 10.1186/s12974-017-0856-z

91 Yoneyama, N., Crabbe, J. C., Ford, M. M., Murillo, A. & Finn, D. A. Voluntary ethanol consumption in 22 inbred mouse strains. Alcohol 42, 149–160 (2008). 10.1016/j.alcohol.2007.12.006

92 Coleman, L. G., Jr. & Crews, F. T. Innate Immune Signaling and Alcohol Use Disorders. Handb Exp Pharmacol 248, 369–396 (2018). 10.1007/164_2018_92

93 Coleman, L. G., Jr., Zou, J., Qin, L. & Crews, F. T. HMGB1/IL-1beta complexes regulate neuroimmune responses in alcoholism. Brain, behavior, and immunity 72, 61–77 (2018). 10.1016/j.bbi.2017.10.027

94 Fraga-Junior, E. B. et al. Attenuation of the levels of pro-inflammatory cytokines prevents depressive-like behavior during ethanol withdrawal in mice. Brain research bulletin 191, 9–19 (2022). 10.1016/j.brainresbull.2022.10.014

95 Schneider, R., Jr., et al. N-acetylcysteine Prevents Alcohol Related Neuroinflammation in Rats. Neurochemical research 42, 2135–2141 (2017). 10.1007/s11064-017-2218-8

96 Vetreno, R. P., Lawrimore, C. J., Rowsey, P. J. & Crews, F. T. Persistent Adult Neuroimmune Activation and Loss of Hippocampal Neurogenesis Following Adolescent Ethanol Exposure: Blockade by Exercise and the Anti-inflammatory Drug Indomethacin. Front Neurosci 12, 200 (2018). 10.3389/fnins.2018.00200

97 Leclercq, S., De Saeger, C., Delzenne, N., de Timary, P. & Starkel, P. Role of inflammatory pathways, blood mononuclear cells, and gut-derived bacterial products in alcohol dependence. Biological psychiatry 76, 725–733 (2014). 10.1016/j.biopsych.2014.02.003

98 Callaghan, C. K. et al. Antidepressant-like effects of 3-carboxamido seco-nalmefene (3CS-nalmefene), a novel opioid receptor modulator, in a rat IFN-alpha-induced depression model. Brain, behavior, and immunity 67, 152–162 (2018). 10.1016/j.bbi.2017.08.016

99 Li, J. et al. Paeoniflorin ameliorates interferon-alpha-induced neuroinflammation and depressive-like behaviors in mice. Oncotarget 8, 8264–8282 (2017). 10.18632/oncotarget.14160

100 Borsini, A. et al. Interferon-Alpha Reduces Human Hippocampal Neurogenesis and Increases Apoptosis via Activation of Distinct STAT1-Dependent Mechanisms. Int J Neuropsychopharmacol 21, 187–200 (2018). 10.1093/ijnp/pyx083

101 Zheng, L. S., Kaneko, N. & Sawamoto, K. Minocycline treatment ameliorates interferon-alpha-induced neurogenic defects and depression-like behaviors in mice. Frontiers in cellular neuroscience 9, 5 (2015). 10.3389/fncel.2015.00005

102 Bi, Q., Shi, L., Yang, P., Wang, J. & Qin, L. Minocycline attenuates interferon-alpha-induced impairments in rat fear extinction. Journal of neuroinflammation 13, 172 (2016). 10.1186/s12974-016-0638-z

103 Zhao, Y. N. et al. Activated microglia are implicated in cognitive deficits, neuronal death, and successful recovery following intermittent ethanol exposure. Behavioural brain research 236, 270–282 (2013). 10.1016/j.bbr.2012.08.052

104 Fernandez-Lizarbe, S., Pascual, M. & Guerri, C. Critical role of TLR4 response in the activation of microglia induced by ethanol. J Immunol 183, 4733–4744 (2009). 10.4049/jimmunol.0803590

105 Fernandez-Lizarbe, S., Montesinos, J. & Guerri, C. Ethanol induces TLR4/TLR2 association, triggering an inflammatory response in microglial cells. J Neurochem 126, 261–273 (2013). 10.1111/jnc.12276

106 Alfonso-Loeches, S., Pascual-Lucas, M., Blanco, A. M., Sanchez-Vera, I. & Guerri, C. Pivotal role of TLR4 receptors in alcohol-induced neuroinflammation and brain damage. J Neurosci 30, 8285–8295 (2010). 10.1523/jneurosci.0976-10.2010

107 Marshall, S. A., Geil, C. R. & Nixon, K. Prior Binge Ethanol Exposure Potentiates the Microglial Response in a Model of Alcohol-Induced Neurodegeneration. Brain Sci 6 (2016). 10.3390/brainsci6020016

108 Carlson, E. R., Melbourne, J. K. & Nixon, K. Pharmacological Depletion of Microglia Protects Against Alcohol-Induced Corticolimbic Neurodegeneration During Intoxication in Male Rats. J Neuroimmune Pharmacol 20, 21 (2025). 10.1007/s11481-025-10173-x

109 Zelinski, E. L., Hong, N. S., Tyndall, A. V., Halsall, B. & McDonald, R. J. Prefrontal cortical contributions during discriminative fear conditioning, extinction, and spontaneous recovery in rats. Exp Brain Res 203, 285–297 (2010). 10.1007/s00221-010-2228-0

110 Mukherjee, S. et al. Alcohol Increases Exosome Release from Microglia to Promote Complement C1q-Induced Cellular Death of Proopiomelanocortin Neurons in the Hypothalamus in a Rat Model of Fetal Alcohol Spectrum Disorders. The Journal of neuroscience : the official journal of the Society for Neuroscience 40, 7965–7979 (2020). 10.1523/JNEUROSCI.0284-20.2020

111 Marshall, S. A. et al. IL-1 receptor signaling in the basolateral amygdala modulates binge-like ethanol consumption in male C57BL/6J mice. Brain, behavior, and immunity 51, 258–267 (2016). 10.1016/j.bbi.2015.09.006

112 Nelson, J. C., Greengrove, E., Nwachukwu, K. N., Grifasi, I. R. & Marshall, S. A. Repetitive binge-like consumption based on the Drinking-in-the-Dark model alters the microglial population in the mouse hippocampus. J Integr Neurosci 20, 933–943 (2021). 10.31083/j.jin2004094

113 Bauer, M. R., McVey, M. M. & Boehm, S. L., 2nd. Three Weeks of Binge Alcohol Drinking Generates Increased Alcohol Front-Loading and Robust Compulsive-Like Alcohol Drinking in Male and Female C57BL/6J Mice. Alcoholism, clinical and experimental research 45, 650–660 (2021). 10.1111/acer.14563

114 Koob, G. F. The dark side of emotion: the addiction perspective. European journal of pharmacology 753, 73–87 (2015). 10.1016/j.ejphar.2014.11.044

115 Elmore, M. R. et al. Colony-stimulating factor 1 receptor signaling is necessary for microglia viability, unmasking a microglia progenitor cell in the adult brain. Neuron 82, 380–397 (2014). 10.1016/j.neuron.2014.02.040

116 Hughes, E. G. & Bergles, D. E. Hidden progenitors replace microglia in the adult brain. Neuron 82, 253–255 (2014). 10.1016/j.neuron.2014.04.010

117 Prakash, A., Zhang, H. & Pandey, S. C. Innate differences in the expression of brain-derived neurotrophic factor in the regions within the extended amygdala between alcohol preferring and nonpreferring rats. Alcoholism, clinical and experimental research 32, 909–920 (2008). 10.1111/j.1530-0277.2008.00650.x

118 Gorka, S. M., Teppen, T., Radoman, M., Phan, K. L. & Pandey, S. C. Human Plasma BDNF Is Associated With Amygdala-Prefrontal Cortex Functional Connectivity and Problem Drinking Behaviors. The international journal of neuropsychopharmacology 23, 1–11 (2020). 10.1093/ijnp/pyz057

119 Fiedler, D. et al. Brain-Derived Neurotrophic Factor/Tropomyosin Receptor Kinase B Signaling Controls Excitability and Long-Term Depression in Oval Nucleus of the BNST. The Journal of neuroscience : the official journal of the Society for Neuroscience 41, 435–445 (2021). 10.1523/jneurosci.1104-20.2020

120 Hsiao, C. C. et al. GPCRomics of Homeostatic and Disease-Associated Human Microglia. Front Immunol 12, 674189 (2021). 10.3389/fimmu.2021.674189

