## Supplementary figures and images for "Microglia promote neurodegeneration and hyperkatifeia during withdrawal and prolonged abstinence from binge alcohol"

### Supplemental Figure 1

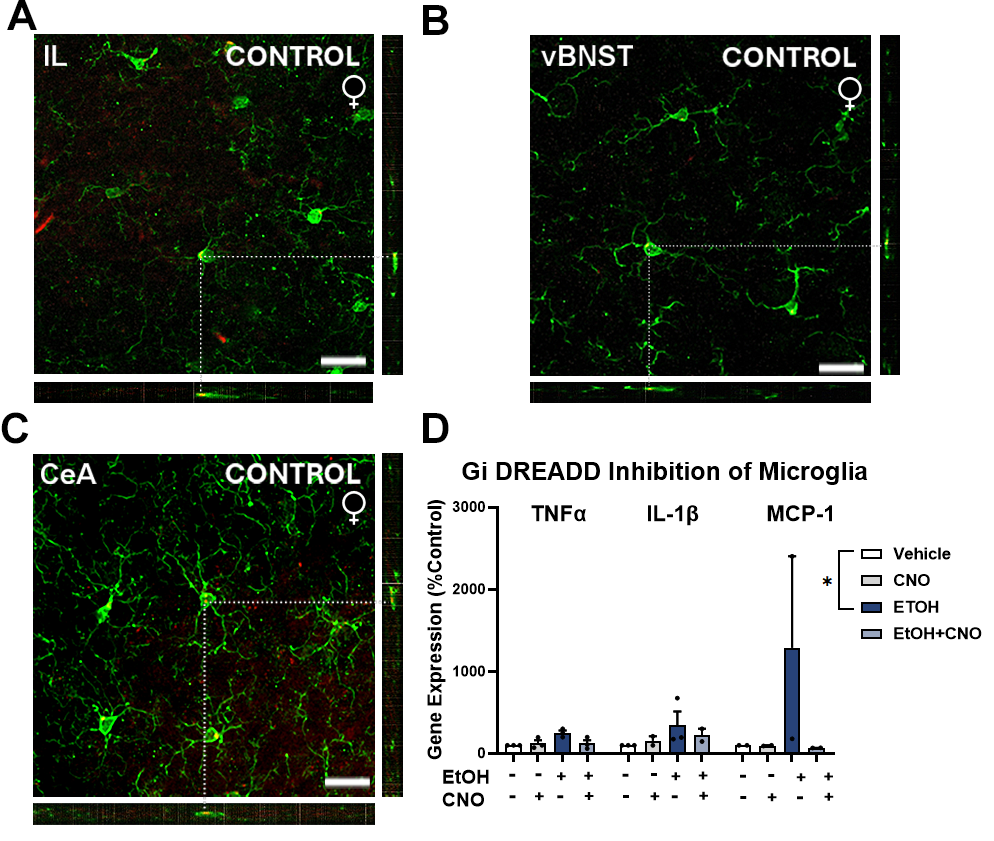

### Supplemental Figure 2

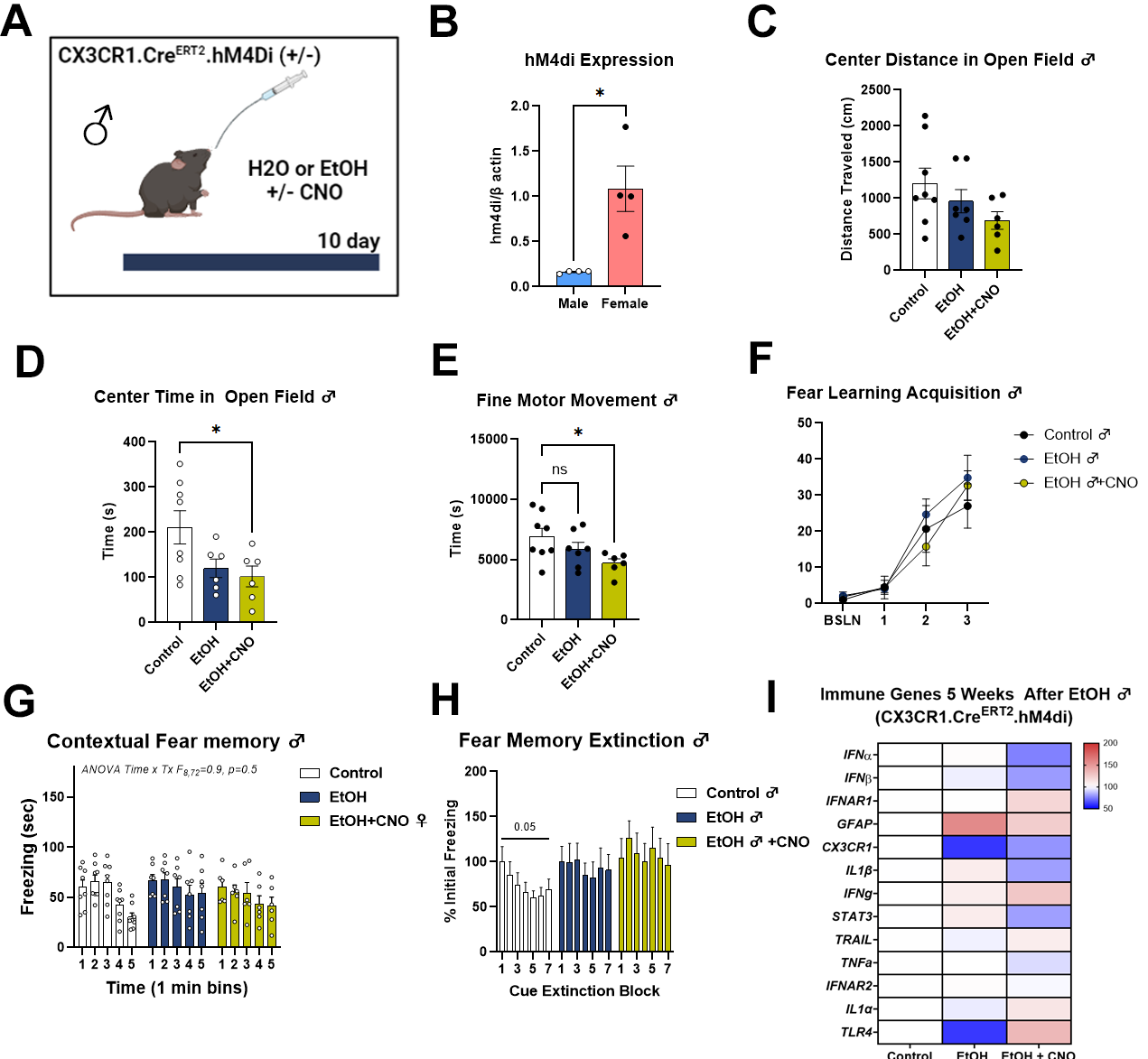
